# Preconditioning of human iPSCs with doxorubicin causes genome-wide transcriptional reprogramming in iPSC-derived cardiomyocytes linked to mitochondrial dysfunction and impaired cardiac regeneration

**DOI:** 10.1101/2025.04.18.649376

**Authors:** Michelle Westerhoff, Nahal Brocke-Ahmadinejad, Yulia Schaumkessel, Karl Köhrer, Arif Dönmez, Roohallah Ghodrat, Andrea Borchardt, Lluís Enjuanes-Ruiz, Julia Tigges, Katharina Koch, Gerhard Fritz, Arun Kumar Kondadi, Andreas S. Reichert

## Abstract

**Background:** The anthracycline doxorubicin (Dox) is a widely used genotoxic chemotherapeutic drug with known dose-limiting cardiotoxic effects. How Dox-induced damage either to cardiomyocytes or to cardiac stem cells, which may compromise cardiac regeneration, contributes to cardiotoxicity remains poorly understood.

**Methods:** Here we used a human induced pluripotent stem cell (iPSC)-based model system to determine the sensitivity of stem cells (iPSCs) and iPSC-derived derived cardiomyocytes (iCMs) applying different treatment regimens of Dox. Next to a broad range of methods to determine cellular and mitochondrial functions we performed an in-depth whole genome transcriptome profiling in iPSCs as well as iCMs.

**Results:** As compared to their differentiated counterparts, iPSCs are highly sensitive against even short pulse-treatments with low Dox concentrations. Using such rather mild treatment conditions, we observed major mitochondrial impairments as demonstrated by increased mitochondrial fragmentation, persistent loss of mitochondrial membrane potential, and reduced ATP levels, while neither a markedly increased nuclear DNA damage response nor apoptosis were detected. Albeit mitochondrial dysfunction was not accompanied by changes in mitochondrial ultrastructure or altered OXPHOS complex assembly, mitochondrial genome (mtDNA) organization was altered. This points to a possible role of mtDNA remodelling for contributing to the high susceptibility of iPSCs to Dox. Whole genome transcriptome profiling revealed major differences in the transcriptional response to Dox treatment between iPSCs and iCMs. We could show that a moderate and transient exposure of iPSCs to Dox is sufficient to cause major transcriptional changes as for example reflected by the downregulation of numerous pivotal genes regulating cellular homeostasis and energy metabolism in iPSCs. Furthermore, pulse-treatment with Dox at the iPSC stage, termed preconditioning here, shifts the global transcriptional landscape of iCMs towards the expression of genes associated with impaired cardiac muscle regeneration, disrupted energy metabolism, altered muscle contraction, and increased fibrosis.

**Conclusions:** Our findings support the hypothesis that Dox-induced mitochondrial dysfunction and transcriptional preconditioning in stem cells results in an impaired regenerative capacity after differentiation. This highlights a potential critical role of stem cells in mediating Dox-induced cardiotoxicity.

## Introduction

The anthracycline doxorubicin (Dox) finds wide applications in the chemotherapeutic treatment of solid tumors (e.g. neuroblastoma, ovary, gastrointestinal, and breast tumors) and hematological malignancies (e.g. lymphatic and myeloid tumors) in adults as well as in children (Sritharan & Sivalingam, 2021). A major clinical problem during, and years after chemotherapy, is the onset of cardiotoxicity resulting from cancer therapy with Dox (Mortensen et al., 1986; Swain et al., 2003). The manifestation of Dox-induced cardiotoxicity depends on the cumulative dose and timing of Dox administration, yet also shows high inter-individual variability in patients (Joerger et al., 2007). Dysregulation of cardiac electrophysiological features within hours to weeks after first administration of Dox has been well documented (Larsen et al., 1992; Steinberg et al., 1987). Side effects of Dox, such as congestive heart failure, left ventricular fraction reduction and serious arrythmias, may occur within the first year of Dox treatment or even after 10 years after the initial application (Mortensen et al., 1986; Mulrooney et al., 2009; Swain et al., 2003). In particular, children, women and elderly patients are prone to late-occurring cardiac problems. The underlying mechanisms of these acute and chronic cardiotoxic effects are still not clear and are a matter of intense research (Avagimyan et al., 2024; Carvalho et al., 2009; McGowan et al., 2017).

This is further complicated by the fact that the regenerative capacity of the heart *per se* and the various proposed mechanisms to counteract cardiac dysfunction are not clear. In recent years, there has been an intensive debate surrounding the possible role of cardiac stem cells for supporting the regenerative capacity of this organ. Concurrently, there are unresolved questions regarding the source of such cells that could potentially differentiate into functional cardiomyocytes in both the embryonic and the adult heart. For decades, the heart was considered as an organ with low regeneration capacity (Karsner et al., 1925). However, upon cardiac failure an increase in cardiomyocyte proliferation has been observed indicating regenerative mechanisms within the heart (Beltrami et al., 2001; Kajstura et al., 1998; Urbanek et al., 2003). In addition, the observed effect of cardiac regeneration has not only been described in cases of cardiac failure, but has also been demonstrated under physiological conditions using ^14^C-labeling approach (Bergmann et al., 2009). Whether this regeneration arises from reactivation of cell division of differentiated cardiac cells or from *de novo* generation of cardiomyocytes from a stem / progenitor cell pool remains unclear. With further investigation of the origin of cardiomyocytes in mammalian heart, a distinct resident population of cells in the heart inheriting stem cell characteristics has been identified and proposed to play a relevant role in the regeneration of heart (Beltrami et al., 2003; van Berlo et al., 2014). Notably, Dox-induced cardiotoxicity is often associated with the induction of fibrosis indicating reduced cardiac regenerative capacity upon Dox treatment (Takemura & Fujiwara, 2007).

In humans, cardiomyocytes represent only 30-35 % of the cellular mass in the heart, yet the homeostasis of these cells was proposed to determine strongly the sensitivity of patients against Dox treatment (Bergmann et al., 2015; Dewing et al., 2022; Litviňuková et al., 2020). Given that cardiomyocytes are terminally differentiated cells, it has been proposed that dysfunction of these cells might not solely be dependent on Dox-induced inhibition of DNA replication (Gewirtz, 1999). Rather the formation of reactive oxygen species (ROS), inhibition of topoisomerases - in particular topoisomerase IIβ in mitochondria-, and inhibition of autophagic processes have been implicated in Dox-induced cardiotoxicity (D. L. Li et al., 2016; S. Zhang et al., 2012).

Besides impairment of cytosolic and nuclear processes, mitochondria have been described to play a pivotal role in the induction of cardiotoxic effects upon Dox exposure (Dhingra et al., 2014; Leboucher et al., 2012; Morris et al., 2023). Given an estimated turnover rate of mitochondrial proteins in the heart of about 16 to 18 days, Dox-induced alterations may persist and propagate over time rather than being eliminated from the pool rapidly (Abdullah et al., 2019; Menzies & Golds, 1971). Yet, in most studies mitochondrial dysfunction has been observed upon treatment with comparably high treatment doses which may not completely reflect the actual clinical situation. In patients, Dox plasma concentrations drop rapidly to 25 nM to 250 nM within 1 hour after short-time infusion of Dox (Choi et al., 2020).

As a consequence, cellular, including mitochondrial, dysfunction observed in *in vitro* experiments administrating high Dox doses likely involves multiple damaging mechanisms (e.g.: nuclear and mitochondrial DNA damage, ROS production, inhibition of autophagic processes, etc.) suggesting the need for more clinically relevant exposure models. We hypothesize that mitochondrial dysfunction could contribute to both long-lasting damage to cardiomyocytes as well as to suppressed cardiac regeneration and that this might play a pivotal role in the manifestation of long-term cardiotoxic effects upon chemotherapeutic treatment of patients with Dox. In sum, it is unclear whether Dox-induced long-term cardiotoxic effects could in part result from impaired regeneration of the heart from cardiac stem cells (CSCs) or from remaining cardiomyocytes, thereby favoring the development of cardiotoxic effects.

Here, we introduce human induced-pluripotent stem cells (iPSCs) and iPSC-derived cardiomyocytes (iCMs) as a cellular model system for CSCs and their differentiated progeny allowing us to dissect whether treatment of iPSCs with rather mild doses of Dox has any consequences in iPSCs and in cells derived from these cells. We show that iPSCs are highly sensitivity to mild and transient Dox treatment regimens. These treatment schemes are insufficient to induce a significant DNA damage response or to impair cardiomyocyte differentiation. However, they are accompanied by substantial and persistent impairments of mitochondrial function in iPSCs. Furthermore, treatment at the iPSC stage causes major alterations in gene expression profiles as well as functional parameters, such as the beating frequency, in cardiomyocytes originating from the early-treated iPSC pool. Our data demonstrated that iPSCs undergo a process of preconditioning that appears to promote fibrosis on one hand, and on the other hand impairs cardiac muscle regeneration pathways. Overall, we propose that the administration of Dox to CSC-like cells during the initial phase of cardiomyocytes differentiation could result in the manifestation of long-term effects impacting cellular remodeling in generated cardiomyocytes. This could represent an underappreciated mechanism contributing to long-term cardiotoxicity after chemotherapy.

## Material and Methods

### Cell lines

Human fetal-derived iPSC line iPS (IMR90)-4 (lentiviral transfection with *Oct4*, *Sox2*, *Nanog* and *Lin28,* WiCell) were used for experimental procedures (Yu et al., 2007). iPSCs were cultured in feeder-free conditions on Geltrex™-coated plates (Thermo Fisher Scientific Inc.) using mTeSR™ Plus media (STEMCELL Technologies Inc.) supplemented with 1 % Penicillin / Streptomycin. Cells were kept under normal oxygen conditions at 37 °C and 5 % CO_2_ with media change every second day. At 80 % confluency, iPSCs were passaged as single cells at a confluency of 80 % using StemPro™ Accutase™ (Thermo Fisher) and a defined number of cells was seeded for the respective experiments. To enhance viability of iPSCs after passaging, media was supplemented with 10 µM ROCK inhibitor Y-27632 2HCl (AdooQ BioScience) for 24 hours followed by media change the next day and subsequent media changes every second day

### Cardiac differentiation

iPSC-derived cardiomyocytes (iCMs) were generated using published protocols (Burridge et al., 2014; Lian et al., 2012). Briefly, 80.000 cells per well were seeded as single cells with StemPro™ Accutase™ (ThermoFisher) on Geltrex™-coated (Thermo Fisher Scientific Inc.) 24-well plates (Greiner Bio-One GmbH) and 10 µM ROCK inhibitor Y-27632 2HCl (AdooQ BioScience). Upon 80 % confluency, cardiac differentiation was initiated by complete aspiration of stem cell media and exchange with basal cardiac differentiation media (CBM) consisting of RPMI 1640 media without L-Glutamine (Capricorn Scientific) supplemented with 1 % Penicillin / Streptomycin (Sigma-Aldrich), 2 mM stable Glutamine (Capricorn Scientific) and 25 mM HEPES buffer (ThermoFisher). During the process of cardiac differentiation, media was completely aspirated and exchanged by the respective cardiac differentiation media. From day 0 to day 2, 1x B-27 supplement without insulin (ThermoFisher), 50 µg / mL ascorbic acid (Sigma-Aldrich) and 6 µM CHIR99021 (MedChemExpress) was added to CBM.

On day 3 of the cardiac differentiation, CBM media was supplemented with 1x B-27 supplement without insulin (ThermoFisher), 50 µg / mL ascorbic acid (Sigma-Aldrich). From day 4 to day 5, 1x B-27 supplement without insulin (ThermoFisher), 50 µg / mL ascorbic acid (Sigma-Aldrich) and 5 µM IWP4 (STEMCELL Technologies Inc.) was added to CBM. From day 5 to day 7 the same media composition was used as described for day 3. From day 7, cells received CBM supplemented with 1x B-27 supplement (ThermoFisher) and 50 µg / mL ascorbic acid (Sigma-Aldrich). Expression of cardiac differentiation markers was checked on the respective days of analysis with quantitative real-time PCR. Primers used are listed in Supplementary Table 1.

### Cell viability assay

To determine the cell viability of iPSCs and iCMs 48 hours after 2 hours pulse-treatment with Dox cells MTT assay (Sigma-Aldrich) was performed. For iPSCs, 6-8×10^4^ cells were seeded as single-cell suspension, on Geltrex™-coated 96-well plates (Greiner Bio-One GmbH). For iCMs, 6×10^4^ cells were split as single-cell suspension on Geltrex™-coated 96-well plates (Greiner Bio-One GmbH). MTT assay measures the reduction of the terazolium salt 3-(4,5-dimethylthiazol-2-yl)-2,5-diphenyltetrazolium bromide by NAD(P)H-dependent oxidoreductase enzymes to purple formazan (Colangelo et al., 1992). Absorbance was measured at 565 nm using Tecan INFINITE 200 Pro (Tecan Austria). Relative cell viability in respective controls was set to 100 %. In addition, cell viability was measured with Sulforhodamine B assay (SRB assay, Supplementary Figure 1). Absorbance was measured using Tecan INFINITE 200 Pro (Tecan Austria). Relative cell viability in respective controls was set to 100 %.

### Gene expression analysis (RT-qPCR)

Total RNA was purified using RNeasy Mini Kit (Qiagen). Reverse transcription was performed with GoScript™ Reverse Transcription Mix, Oligo(dT) kit (Promega) using 1 µg RNA. qRT-PCR was performed as followed: 1.25 °C – 5 min; 2. Extension at 42 °C – 60 min; 3. Inactivation with 70 °C – 15 min and holding of the samples at 4 °C. Quantitative analyses were performed in triplicates using Corbett Rotorgene RG-6000 (Corbett Research) and the GoTaq® qPCR Master Mix (Promega). Quantitative real-time PCR (qRT-PCR) was performed as followed: 1. 95°C – 2min; 2. Amplification for 40 cycles with 95°C - 5s, 60°C - 10 s; 3. Melting curve analysis with 1 °C degree steps ramping from 65 °C to 95 °C. Primers for qRT-PCR are listed in Supplementary Table 1. At the end of each run, melting curves were analyzed. PCR products with Cq-values ≥ 35 were neglected. HPRT1 was used as housekeeping gene. For mtDNA:gDNA ratio, GAPDH was used as housekeeping gene. Relative mRNA expression was set to 1.0 for the untreated controls.

### Immunofluorescent staining of cells

To perform immunofluorescence staining, 7.5 x10^4^ cells per well were seeded as single-cells on Geltrex™-coated cover glasses (Carl Roth) of a 6-well plate (Greiner Bio-One GmbH). Briefly, at the day of analysis cells were fixed with 4 % PFA for 15 min at room temperature, followed by second fixation with 100 % methanol for 10 min at −20 °C. Samples were permeabilized and blocked in PBS with 0.5 % Triton X-100, 5 % BSA (PAN-Biotech) for 30 min at room temperature, and then incubated with primary antibody in PBS with 0.1 % Triton X-100, 5 % BSA over night at 4 °C as indicated. Samples were then washed, incubated with secondary antibody for 1 h at room temperature, and mounted with ROTI®Mount FluorCare DAPI (Carl Roth). Following antibodies were used: α-Ki-67 (Cell Signaling, #9449S), α-γH2AX (Cell Signaling, # 80312S), α-53BP1 (Cell Signaling, # 4937), α-HSP60 (Sigma Aldrich®, # SAB4501464-100UG), α-dsDNA (PROGEN, #AC-30-10), goat anti-mouse IgG H&L (HRP) (abcam, #ab6789), goat anti-rabbit IgG H&L (HRP) (BIOZOL, # 111-035-144). Fluorescence Z-stack imaging of iPSCs was acquired at ECLIPSE Ti2 inverted confocal microscope (Nikon Instruments Inc.), coupled with UltraVIEW®VoX spinning disc laser system (PerkinElmer Inc.) equipped with a 63-x oil objective (N.A. 1.2) using Velocity 6.3 software (Perkin Elmer Inc.).

### SDS PAGE and Western Blotting

For SDS-PAGE, at least 7.5 x10^4^ cells per well were seeded onto Geltrex™-coated 6-well plates (Greiner Bio-One GmbH). After following the Dox treatment regimen including a recovery time of 48 hours, cells were harvested and pellets were frozen at −80 °C. Pellets were thawed on ice and protein was extracted with RIPA lysis buffer. Protein levels were quantified using *DC* Protein Assay Kit (Bio-Rad Laboratories, Inc.). Protein lysates (15 µg) were separated on 15 % or 5 % SDS-gels, transferred to nitrocellulose membranes (Amersham) and decorated with antibodies as indicated. Signal was visualized with Thermo Scientific™ SuperSignal™ West Femto Maximum Sensitivity Substrate (Thermo Fisher Scientific Inc.) and Fusion SL (Vilber). Intensity of the bands were analyzed using ImageJ 1.53c (Rasband, 2020) and normalized to untreated control. Following antibodies were used: α-PARP (Cell Signaling, #953S), α-Caspase3 (abcam, #ab32351), α-cleaved Caspase 3 (Cell Signaling #9664S), α-Actin (Thermo Fisher Scientific Inc., #MA1-744), α-H2AX (Sigma Aldrich®, #SAB5600038-100UG), α-ATM (Cell Signaling, #2873T), α-pATM (Thermo Fisher Scientific Inc., MA1-46069), α-Vinculin (Sigma Aldrich®, V9131-100UL), α-OPA1 (Pineda, polyclonal rabbit antibody custom-made against C-terminal peptide (Duvezin-Caubet et al., 2006)), goat anti-mouse IgG H&L (HRP) (abcam, #ab6789), goat anti-rabbit IgG H&L (HRP) (BIOZOL, # 111-035-144).

### Analysis of apoptosis and DNA damage response

For measurement of apoptosis in iPSCs, 7.5×10^4^ cells were seeded as single cells onto Geltrex™-coated 6-well plates (Greiner Bio-One GmbH). The pan caspace inhibitor Z-VAD-FMK (AdooQ BioScience) was used in order to limit cell death. Cells were pre-incubated with Z-VAD-FMK for 4 hours prior to procedure of Dox treatment regimen. Staurosporine was applied as a positive control. Induction of apoptosis was determined using SDS-PAGE and western blot analysis using indicated markers. Protein levels of each treatment were determined and further evaluated using the software GrapPad Prism8. To assess DNA damage induced by moderate pulse-treatment with Dox, the levels of DNA damage markers phosphorylated histone variant H2A.X (γH2AX, phosphorylated histone H2A.X at serine 139) and P53-binding protein 1 (53BP1) were measured by immunofluorescence staining. Briefly, 2.5×10^4^ cells (iPSCs) per wells were seeded on Geltrex™-coated coverslips in 24-well plate Greiner Bio-One GmbH) followed by treatment regimen with Dox. Here, 10nM of etoposide (Eto) treatment was used as a positive control. Cells were then fixed at 0.5, 4-, 8-, and 24-hours post-treatment and stained for the respective markers. Subsequent to this, the number of individual γH2AX and 53BP1 foci, as well as colocalizing foci per nucleus, was manually counted using ImageJ (version 1.53c). The average number of DNA damage foci per nucleus was calculated, and further data analysis was performed using GraphPad Prism 8.

### MitoTracker™Green FM and TMRM staining

For analysis of mitochondrial morphology and membrane potential, 7.5×10^4^ cells were seeded on Geltrex™-coated 3.5 cm MatTek glass bottom dishes (MATTEK) for live-cell microscopy followed by treatment regimen with Dox. On the day of microscopy, cells were stained with MitoTracker™Green FM and Tetramethylrhodamin (TMRM) for 30 min (Thermo Fisher Scientific Inc.). Cells were washed thrice with 1 x PBS and were covered 1 mL mTeSR™ Plus (STEMCELL Technologies Inc.) and 10 µM HEPES buffer was added for buffering oxidation of media. Fluorescence imaging of iPSCs was ECLIPSE Ti2 inverted confocal microscope (Nikon Instruments Inc.), coupled with UltraVIEW®VoX spinning disc laser system (PerkinElmer Inc.) equipped with a 63-x oil objective (N.A. 1.2) using Velocity 6.3 software (Perkin Elmer Inc.). Z-stack imaging was performed at 37 °C. For analysis of mitochondrial morphology with MitoTracker™Green FM, images were randomized and mitochondrial morphology was categorized as tubular, intermediate or fragmented. For estimation of mitochondrial membrane potential, in Velocity 6.3 software (Perkin Elmer Inc.) for each image the background was manually corrected and for each individual cell a region of interest (ROI) was manually defined prior to measurement of TMRM fluorescence intensity. Outliers were identified and removed using ROUT (Q=1) outliers test in GraphPad Prism 8 (GraphPadPrism Software, Inc.). Relative TMRM fluorescence intensity for the untreated control was set to 1.0.

### Analysis of cellular bioenergetics and energy metabolism

Optimal seeding density and FCCP concentration without toxicity response were tested before starting the Seahorse XF Cell Mito Stress Test assay (Agilent Technologies, Inc). For Seahorse run, iPSCs were pulse-treated with Dox for 2 hours followed by 24 hours of recovery. Afterward, 3.5 x 10^3^ cells per well were seeded on Geltrex™-coated 96-well Seahorse XF cell culture plates and further recovery of 24 hours was continued. According to manufacturer’s protocol, XF sensor cartridges were hydrated with Seahorse XF Calibrant at 37 °C in a non-CO_2_ incubator overnight. Normal cell culture medium was changed to phenol-free Seahorse XF DMEM (Agilent Technologies, Inc) containing 10 mM Glucose, 1 mM Pyruvate and 2 mM L-Glutamine (all from Sigma Aldrich®). OCR was measured in a Seahorse XFe96 Flux Analyzer with Seahorse Wave 2.4 software (Agilent Technologies, Inc). In 3 cycles of 3 min mixing and 3 min recording, 1 µM Oligomycin, 0.125 µM FCCP and 0.5 µM Antimycin/Rotenone AA was injected and OCR was measured. The data was normalized to the cell number using Hoechst normalization assay (Thermo Fisher Scientific Inc.). Relative OCR (pmol/min/cell) was set to 100 % for the control samples.

### Analysis of ATP levels

To determine ATP levels in treated and untreated iPSC, 8×10^3^ cells were seeded onto Geltrex™-coated 96-well glass bottom plates (Greiner Bio-One GmbH). After recovery time, luminescence-based assay CellTiter-Glo® 2.0 Cell Viability Assay (Promega) was performed. Luminescence was measured at CLARIOStar^Plus^ plate reader (BMG Labtech GmbH). The data was normalized to the cell number using the Hoechst normalization assay (Thermo Fisher Scientific Inc.). Relative ATP levels of control cells was set to 1.0.

### Analysis of mitochondrial DNA organization

For mitochondrial nucleoid imaging, Z-stack images (20 stacks per image) were recorded using an ECLIPSE Ti2 inverted confocal microscope (Nikon Instruments Inc.), coupled with an UltraVIEW®VoX spinning disc laser system (PerkinElmer Inc.) equipped with a 63-x oil objective (N.A. 1.2) using Velocity 6.3 software (Perkin Elmer Inc.). Using ImageJ 1.53c [3], each z-stack image was projected to one z-plane (Maximum projection) followed by separation of each channel and manual thresholding of 8-bit images (Otsu-Threshold). Watershed function was performed and images were further proceeded using a working pipeline in CellProfiler4.2.5 (Stirling et al., 2021). Cells were analyzed with a semi-automated pipeline in CellProfiler. The number of mitochondrial nucleoids per areas of cell and the size of mitochondrial nucleoids (µm^2^) was determined.

### Electron microscopy

For investigation of the ultrastructure of mitochondria, 7.5×10^4^ cells per well were seeded into Geltrex™-coated 6-well plates (Greiner Bio-One GmbH). iPSCs were fixed with 3 % glutaraldehyde / 0.1 M sodium cacodylate buffer at pH 7.2. Cell pellets were washed in fresh 0.1 M cacodylate buffer at pH 7.2 and embedded in 3 % low melting agarose. Staining and filtration was performed in 1 % osmium tetroxide for 50 min, two washing steps with cacodylate buffer and once with 70 % EtOH for 10 min each. Further staining was proceeded in 1 % uranyl acetate / 1 % phosphotunstic acid in 70 % EtOH for 1 h. Dehydration of samples was performed by incubation of samples once in 90 % EtOH, once in 96 % EtOH, thrice in 100 % EtOH, and lastly twice in 100 % acetone. Stained samples were embedded in low viscosity EMS for polymerization at 70 °C for 48 hours. Ultrathin sections (70 nm) of each sample were prepared using a Leica EM UC7microtome (Leica Biosystems GmbH) images were acquired with a JEM-2100 Plus transmission electron microscope (JEOL, Freising, Germany) at 200 KV equipped with an EM-24830 flash CMOS camera system (JEOL, Freising, Germany). Images of ultrastructure were randomized and were categorized into tubular, septae, arched and branched cristae, and the mean percentage of each cristae phenotype was determined.

### Blue Native and Clear Native PAGE

For Blue Native and Clear Native PAGEs, 7.5×10^4^ cells per well were seeded into Geltrex™-coated 6-well plates (Greiner Bio-One GmbH). iPSCs were treated according to the defined treatment scheme (see Fig. 1A) and harvested after 48 hours of recovery time. Cell pellets were resuspended in 1 ml lysis buffer (210 mM mannitol, 70 mM sucrose, 1 mM EDTA, 20 mM HEPES, 0.1 % BSA, 1 x protease inhibitor) and incubated on ice for 10 min. Cells were disrupted using a 20G canula, followed by sequential centrifugation steps at 1000 x g, 4 °C for 10 min to remove cell debris and 10,000 x g, 4 °C for 15 min to pellet mitochondria. Mitochondrial pellet was resuspended in BSA-free lysis buffer and protein concentration was determined using DC Protein Assay Kit.

**Figure 1.**
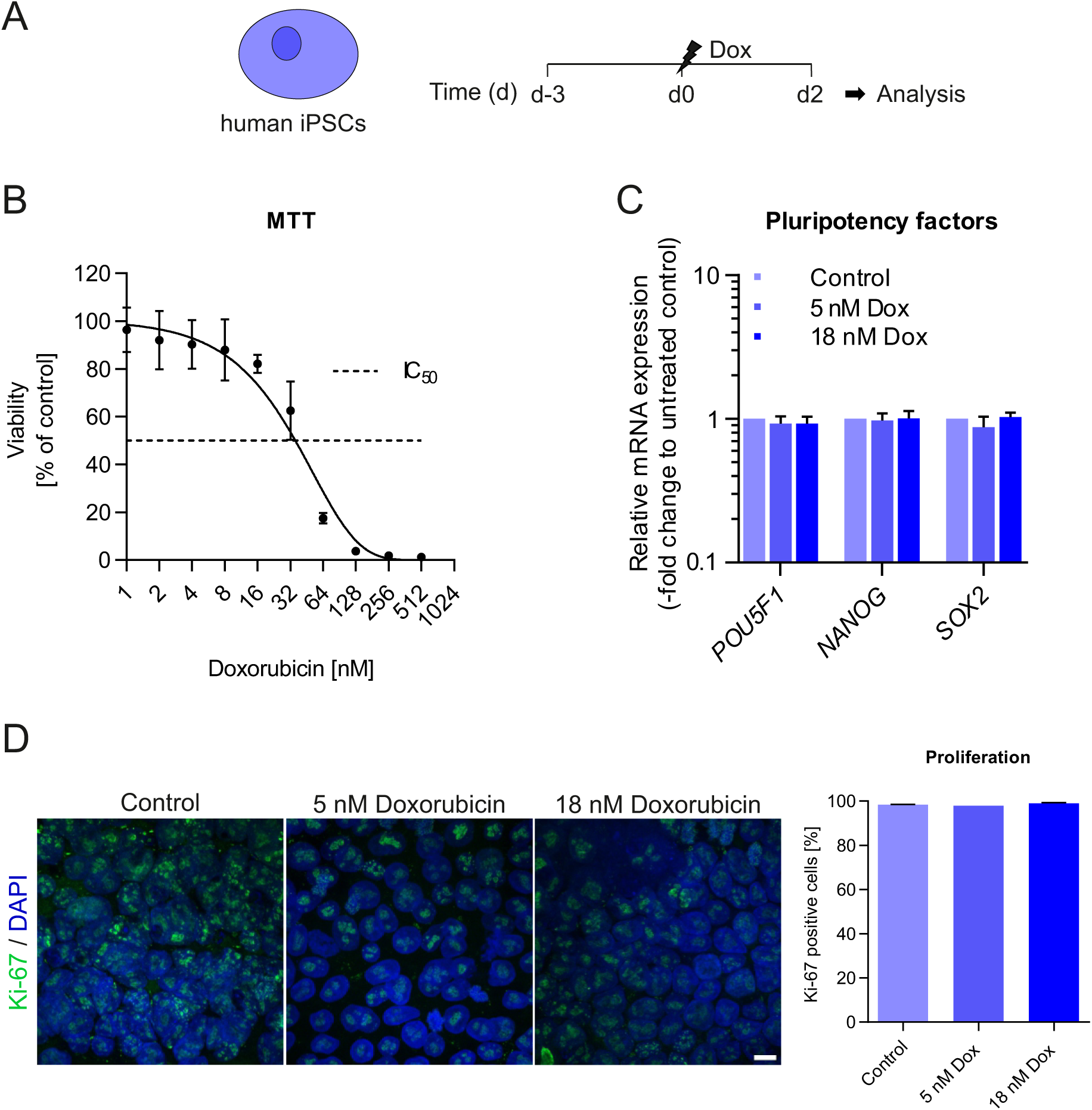
Human induced-pluripotent stem cells (iPSCs) are highly sensitive to doxorubicin pulse-treatment. **A** Schematic illustration of pulse-treatment scheme with Dox in iPSCs. Pulse-treatment with Dox is represented by a lightning symbol. **B** Viability of iPSCs was elucidated 48 hours after pulse-treatment with Dox using MTT assay. Data points represent the mean ± SEM of three independent experiments (n=3; N=8). Dashed line marks the IC_50_-value. **C** Quantitative real-time PCR results of the mRNA expression of stem cell markers POU Class 5 Homeobox 1 (*POU5F1*), Nanog Homeobox (*NANOG*) and SRY-Box Transcription Factor 2 (*SOX2*). Relative mRNA expression of Dox-treated iPSC compared to the control represent the mean ± SEM (n=5; N=3). **D** Immunohistochemical detection of the proliferation marker Ki-67 (green color) in Dox-treated and untreated iPSC. DNA was counterstained with DAPI (blue color). Left side, representative images. Magnification: 63x, scalebar 10 µm. Right side, quantitative results of two independent experiments of Ki-67 positive cells in iPSC 48 h after Dox treatment. Data represents the average percentage ± SEM of two independent experiments (n=2; N =100 nuclei).

For blue native page, 100 µg of mitochondria were solubilized for 1 h on ice using 2.5 g / g of digitonin to protein ratio. Insolubilized material was pelleted at 20,000 x g, 4 °C for 20 min. The supernatants were supplemented with loading buffer (50 % glycerol, 8 g / g Coomassie to detergent ratio) and loaded onto 3-13 % gradient gel and run at 150 V, 15 mA, 4 °C for 16 h. Thereafter, protein complexes were transferred onto PVDF membrane and blocked overnight with 5 % milk in TBS-T at 4 °C. For identification of relevant protein complexes, the membrane was decorated using the following antibodies: NDUFB4 (abcam, #ab110243), UQCRC2 (abcam, #ab14745), COXIV (abcam, #ab16056) ATP5A (abcam, #ab14748), OXPHOS cocktail antibody (abcam, #ab110412, 1:2000), Mic60 (abcam, #ab110329), Goat IgG anti-Mouse IgG (abcam, #ab97023) and Goat IgG anti-Rabbit IgG (Dianova, #SBA-4050-05) conjugated to HRP. The chemiluminescent signals were obtained using Pierce™ SuperSignal™ West Pico PLUS chemiluminescent substrate reagent (Thermo Scientific) and VILBER LOURMAT Fusion SL equipment (Peqlab).

For clear native gels, 300 µg mitochondria were solubilized on ice for 1 h with 2.5 g / g digitonin to protein ratio. The samples were centrifuged for 20 min at 20,000 x g and 4 °C to pellet insolubilized material. The supernatants were supplemented with loading buffer (50 % glycerol, 0.01 % Ponceau S) and immediately loaded onto 3-13 % gradient gel. Complexes were separated at 150 V, 15 mA for 16 h. To assess complex in-gel activity, the gel slices were incubated in respective buffer solution for several hours at room temperature. For complex I activity, the gel was incubated in 5 mM Tris-HCl pH 7.4, 0.1 mg / mL NADH and 2.5 mg / mL nitro blue tetrazolium chloride (NBT). For complex IV activity, the gel was incubated in 50 mM sodium phosphate buffer pH 7.2, 0.05 % DAB and 50 µM horse heart cytochrome *c*.

### Quantification of mitochondrial DNA (mtDNA) copy number

For this approach, 7.5×10^4^ cells per well were seeded into Geltrex™-coated 6-well plates (Greiner Bio-One GmbH). Analysis of mitochondrial DNA content in relation to genomic DNA content was examined using qPCR. For each sample, total DNA was isolated using DNeasy Blood & Tissue Kit (QIAGEN). Analyses were performed using Corbett Rotorgene RG-6000 (Corbett Research). Quantitative PCR was performed as followed: 1. 95°C – 2min; 2. Amplification for 40 cycles with 95°C - 5s, 60°C – 10 s; 3. Melting curve analysis with 1 °C degree steps ramping from 65 °C to 95 °C. Primers are listed in Supplementary Table 1. The Cq-values of mitochondrial genes Mitochondrially Encoded NADH:Ubiquinone Oxidoreductase Core Subunit 1 (*MT-ND1*) and RNA, Ribosomal 45S Cluster 2 (*RNR2*) were normalized to genomic gene Glyceraldehyde-3-Phosphate Dehydrogenase (*GAPDH*). Relative mtDNA copy number of the control was set to 1.0.

### Beating characterization of human induced-pluripotent stem cell derived cardiomyocytes (iCMs)

For beating analysis, 8×10^4^ cells per well were seeded into Geltrex™-coated 12-well plates (Greiner Bio-One GmbH). On day 9 of iCM differentiation, live cell videos of about 20 seconds were acquired with x20 magnification at Leica DM IL LED Fluo Cellfactory (Leica) using 60 frames per second. All video analyses of beating frequency in the wells was conducted using an automated system applied earlier (Galanjuk et al., 2022). The following steps were applied to analyze each video. First, a motion profile was established by placing reference points on the video in a grid pattern, with each point spaced 20 pixels apart. The motion of each reference point was then tracked using the Lucas-Kanade method (Lucas & Kanade, 1981), which is available in the OpenCV library (Bradski, 2000). This tracking was conducted on a frame-by-frame basis. The movement of each reference point generated a distance profile, indicating how far each point moved from its initial position in the first frame at time t. Next, a Savitzky-Golay filter (Savitzky & Golay, 1964) was applied to the resulting time-distance curves (motion profiles) to filter out minimal movements caused by noise or potential artifacts like floating particles. This filtering is crucial for distinguishing between the motion of cells and any artifacts, helping to reduce noise in the analysis. Only the motion profiles of grid points within a manually pre-defined area characterized by clearly visible contractions were considered for further analysis. By performing peak analysis on the remaining distance profile curves of the grid points, the endpoint beating frequency was determined. Exemplary distance profiles are shown in the supplementary material (Supplementary Figure 8).

### Whole genome transcriptome analysis

For the whole genome transcriptome approach, 8×10^4^ iPSCs per wellwere seeded with StemPro™ Accutase™ (ThermoFisher) on Geltrex™-coated 12-well plates (Thermo Fisher Scientific Inc.) and 10 µM ROCK inhibitor Y-27632 2HCl (AdooQ BioScience) for 24 hours. When iPSCs reached 80 % confluency, cardiomyocyte differentiation was initiated by following a chemically defined protocol. Following treatment conditions were included: iPSC untreated (iPSC_Mt), iPSC treated with Dox (iPSC_Dox), untreated iPSC-derived cardiomyocytes (iCM_Mt_Mt), iCMs treated at iPSC stage (iCM_Dox_Mt), iCMs treated on d7 of cardiac differentiation (iCM_Mt_Dox) and iCMs treated at iPSC and iCM stage (iCM_Dox_Dox). For iPSCs, cells were harvested 48 hours after 2 hours pulse-treatment with Dox (respective to IC_30_-value). For iCMs, cells were treated with Dox for 2 hours (IC_30_-value respective to cell type) and all iCM samples were harvested on day 9 of cardiac differentiation. For preparation of transcriptomic samples, total RNA was purified using RNeasy Mini Kit (Qiagen). The expression for pluripotent factors POU Class 5 Homeobox 1 (*POU5F1*), Nanog Homeobox (*NANOG*) and cardiac markers Troponin T2 (*TNNT2*), NK2 Homeobox 5 (*NKX2-5*), Myosin Heavy Chain 6 (*Myh6*) and GATA Binding Protein 4 (*GATA4*) was determined with qRT-PCR before further procedure. For each sample the RNA concentration was determined and 20 ng RNA was aliquoted for transcriptomics analysis. All RNA samples were quantified (Qubit RNA HS Assay, Thermo Fisher Scientific, MA, USA) and quality was assessed by capillary electrophoresis using the Fragment Analyzer and the ‘Total RNA Standard Sensitivity Assay’ (Agilent Technologies, Inc. Santa Clara, CA, USA). Following manufacturer’s protocol, library preparation was performed using ‘VAHTS™ Stranded mRNA-Seq Library Prep Kit’ for Illumina®. As input for mRNA capturing, fragmentation, cDNA synthesis, adapter ligation and library amplification, 20 ng of RNA was used. The bread purified library was normalized and sequencing was performed on the NextSeq2000 system (Illumina Inc. San Diego, CA, USA) with a read setup of 1×100 bp. Data was converted from bcl files to fastq files using the BCL Convert Tool (version 3.8.4). Additionally, adapter trimming and demultiplexing was performed with this tool. Quality control was done and submitted together with the read fastq files. The stranded library of single end mRNA reads was mapped to the human reference genome (ENSEMBL, GRCh38) using the tool HISAT2 (Kim et al., 2019). Alignments were sorted with samtools (H. Li et al., 2009) sort option and counts per transcript were received from applying HTSeq (Anders et al., 2015). The subsequent analysis was performed in R (version 4.3.0) using the Bioconductor (Huber et al., 2015) packages edgeR (Robinson et al., 2009) and limma/voom (Law et al., 2014; Ritchie et al., 2015) to create a count matrix, to filter low expression profiles and to identify differentially expressed genes. Based on the result of multidimensional scaling (Ritchie et al., 2015) which reveals the relationships between the expression profiles of the samples, one replicate was discarded from further analysis to avoid biased results (Replicate number 4 of Sample iPSC_Dox, Supplementary Figure 4). Before testing for differential expression, the expression based on the count data was normalized by trimmed mean of M values (TMM, (Robinson & Oshlack, 2010)) and counts were transformed to lcpm (log counts per million reads). Pairwise comparisons lead to sets of significant differentially expressed genes (Benjamini Hochberg adjusted p-value < 0.05) which were further analyzed by GO-term enrichment using the R packages ClusterProfiler (Wu et al., 2021). Heatmaps were generated using the Bioconducter package ComplexHeatmaps (Gu, 2022; Gu et al., 2016).

### Statistical analysis

Data is represented as mean ± standard error of mean (SEM). Statistical significance between two groups was determined using unpaired Student’s t-test. Statistical significance between more than two groups was determined by one-way ANOVA followed by Dunnett’s or Turkey’s *post hoc* test. *P-value ≤ 0.05, **P-value ≤ 0.01, ***P-value ≤ 0.001, ****P-value ≤ 0.0001. Data analysis was performed using Microsoft Excel. Data representation and statistical analysis was performed using GraphPad Prism8.

## Results

### Human iPSCs are highly sensitivity to pulse-treatment with doxorubicin

Plasma concentrations of Dox in cancer patients receiving fast infusions have been observed to decline rapidly reaching low steady-state concentrations between 25 nM and 250 nM (Gewirtz, 1999). Previously, we established an *in vitro* treatment scheme aiming to mimic the clinical situation in patients using a pulse-treatment with Dox comprising 2 hours treatment with concentrations ranging from 10 nM to 500 nM (Jahn et al., 2020). Here, we treated iPSCs for 2 hours with Dox followed by a 48 hours recovery phase (Fig. 1A). To elucidate the toxicity of Dox on iPSCs under these conditions, cell viability of iPSCs with increasing Dox concentrations (range 1 nM to 512 nM) was determined 48 hours after Dox pulse-treatment using an MTT assay (Fig. 1B). iPSCs showed a remarkably high sensitivity to Dox (IC_50_-value = 35 nM) despite the short exposure time of 2 hours. This was corroborated using another assay commonly used, namely Sulforhodamine B assay (Supplementary Figure 1). Due to the high sensitivity of iPSCs and the rationale to ideally induce stress only moderately, we used doses within the IC_10_ and IC_30_-values (5 nM and 18 nM, respectively) for all subsequent experiments. In this way, only mild and sub-lethal stress were applied which still would enable the differentiation of Dox-treated iPSCs into cardiomyocytes. To ascertain whether iPSCs maintain their pluripotency when exposed to moderate doses of Dox, expression levels of established pluripotency markers *POU5F1*, *NANOG* and *SOX2* were determined. No apparent differences in the expression of pluripotency markers were detectable (Fig. 1C) demonstrating that the pluripotent state of iPSCs was not affected upon pulse-treatment with Dox. To analyze whether the proliferation capacity of iPSCs was maintained upon Dox treatment, the extent of Ki-67, a well-known proliferation marker, detectable within the cell nucleus was determined via immunofluorescence staining. Dox treatment did not cause any reduction in this marker (Fig. 1D) demonstrating that proliferation of iPSCs was not significantly altered upon Dox treatment. Thus, short pulse-treatment of iPSCs with the chosen sub-lethal doses of Dox did not result in a detectable change in the expression of surrogate markers for pluripotency and proliferation of iPSCs.

### Mild pulse-treatment with doxorubicin is not sufficient to induce a major DNA damage response or apoptosis

The intricate relation between DNA damage repair (DDR) and apoptosis induction upon genotoxic stress is well known (Wei et al., 2006). For stem cells exposed to genotoxic substances, DDR and regulation of apoptosis are essential for survival and stem cell pool maintenance. To analyze whether the mild and short Dox treatment conditions result in the induction of a DDR, several markers of this pathway were determined over time. First, the DDR marker proteins Ser139-phosphorylated histone variant H2AX (γH2AX) and tumor suppressor p53-binding protein 1 (53BP1) were quantified for the first 24 hours using immunofluorescence staining. Etoposide (Eto) was used as a positive control for the induction of DNA double-strand breaks (DSBs). Compared to the control, neither of these two markers exhibited a discernible increase overtime following Dox treatment. In contrast, the administration of Eto resulted in a significant increase (Fig. 2A). To further validate that DDR is not activated upon transient exposures of iPSCs to moderate doses of Dox, protein expression levels of γH2AX, protein kinase ataxia telangiectasia mutated (ATM), and its activated phosphorylated form pATM were determined over the same time-course using immunoblotting. Also here, neither of these markers showed a Dox-dependent increase as compared to the control (Fig. 2B,C). We conclude that Dox as applied here did not induce DSBs to a major extent suggesting that the observed high sensitivity of iPSCs towards Dox is possibly not caused by excessive induction of DSBs in the nucleus.

**Figure 2.**
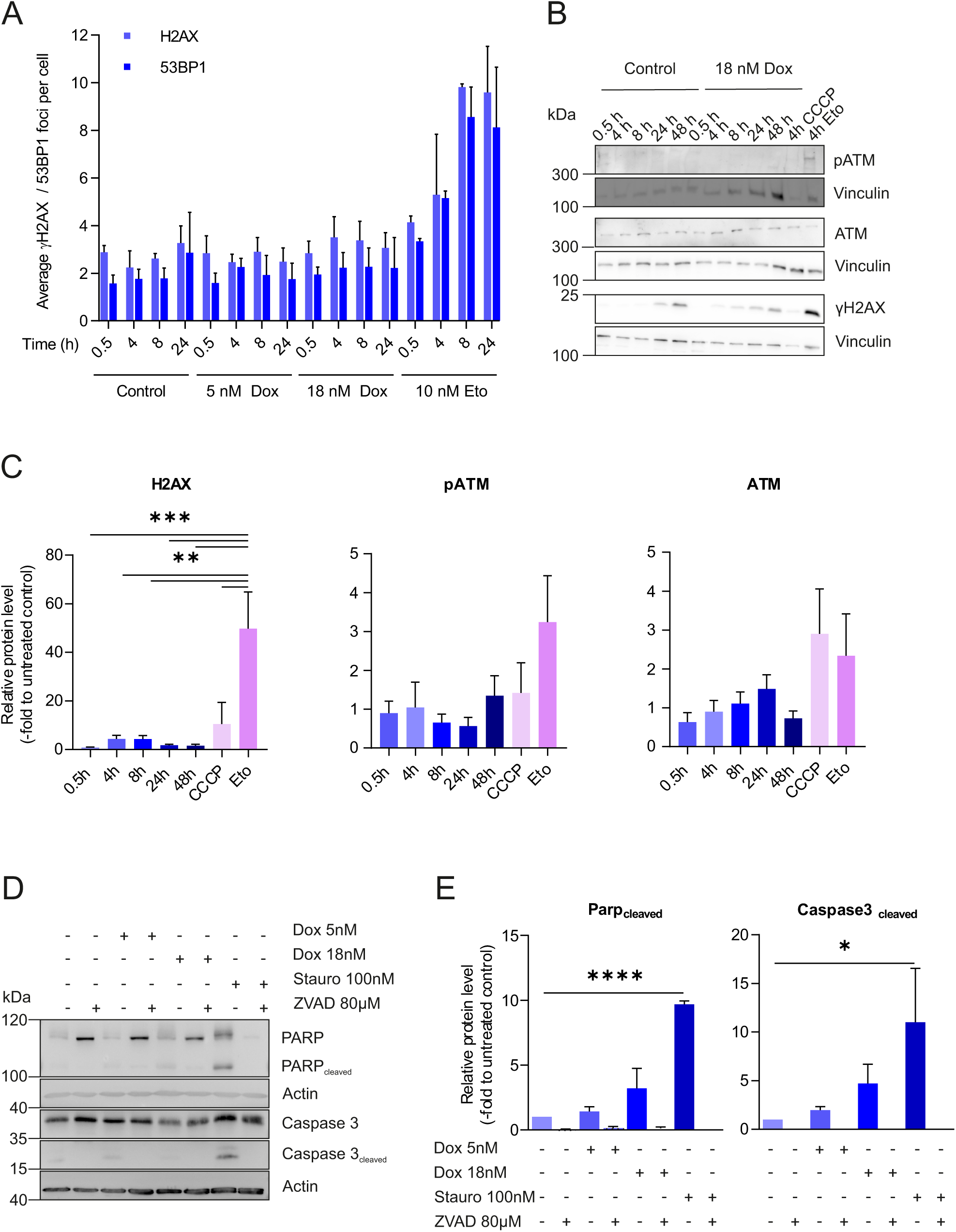
Doxorubicin pulse-treatment is not accompanied with DNA damage response or apoptosis in human iPSCs. **A** Quantitative analysis of immunohistochemical detection of the DNA damage marker γH2AX and 53BP1 0.5, 4, 8 and 24 hours after 2 hours Dox pulse-treatment. Data points shown are the mean ± SEM (n=3; N=30-100 nuclei). As positive control, iPSCs were treated with 10 nM Etoposide (Eto) for 2 hours. **B** Representative immunoblots of DNA damage markers γH2AX, ATM, and its activated phosphorylated form pATM (Ser1981). Treatment conditions and timepoints of analysis are indicated. 10 µM Carbonylcyanid-m-chlorphenylhydrazon (CCCP) and 10 nM etoposide (Eto), each for 4 hours, served as positive controls for mitochondrial and genomic stress, respectively. **C** Quantification of relative protein expression levels of DNA damage marker proteins in Dox-treated iPSCs compared to their respective timepoint control. For CCCP and Eto, relative expression levels were normalized to the 4 h control timepoint. Shown are relative protein expression levels ± SEM (n=3). Vinculin was used as loading control. **D** Representative western blots of apoptosis markers PARP (total and cleaved PARP), Caspase 3, and Caspase 3-cleaved products was determined 48 hours after treatment. **E** Relative protein levels of untreated iPSC were set to 1.0. Actin was used as loading control. Quantitative data represents the mean ± SEM (n=3). Quantification was performed with ImageJ software. For statistical analysis, One_-_way ANOVA with multiple comparisons was performed. * p-value ≤ 0.05, ** p-value ≤ 0.01, *** p-value ≤ 0.001, **** p-value ≤ 0.0001.

To elucidate whether Dox exposure induces apoptosis in iPSCs, protein levels of established markers involved in the apoptotic cascade were determined using immunoblotting. Short pulse-treatment of iPSCs with Dox was not sufficient to induce PARP cleavage and Caspase 3 activation in a significant manner, only a moderate non-significant increase was observed using 18 nM Dox (Fig. 2D,E). As expected, treatment with the positive control staurosporine caused a significant induction of apoptosis. Consistent with our results on DDR, we conclude that pulse-treatment of iPSCs with moderate doses of Dox does neither induce DDR nor apoptosis to a major extent. These findings distinguish this treatment scheme from others in the literature using much higher Dox concentrations and/or incubation times.

### Doxorubicin induced mitochondrial fragmentation in iPSCs is accompanied with loss of mitochondrial membrane potential

In order to check whether short-pulse-treatment of iPSCs with low doses of Dox is accompanied with changes in mitochondrial function, mitochondrial morphology was studied using live-cell confocal microscopy in iPSCs and mitochondrial morphology was categorized into one of the following three categories: ‘tubular’, intermediate’ or ‘fragmented’. Overall, the average percentage of fragmented mitochondria was significantly elevated already at the lowest concentration of Dox of 5 nM (Fig. 3A,B). Consistent with this, we observed a moderate, yet significant, increase of OPA1 processing (Supplementary Figure 2) with 18 nM DOX indicating an impairment of mitochondrial functionality even upon short pulse-exposures with moderate doses of Dox. To elucidate whether Dox pulse-treatment is sufficient to induce an imbalance in mitochondrial membrane potential (ΔΨ) in iPSCs we measured the relative intensity of TMRM within mitochondria using live-cell confocal microscopy. Here, even after a recovery period of 48 hours Dox pulse-treatment was sufficient to significantly reduce ΔΨ of mitochondria in iPSCs in a persistent manner (Fig. 3C, D). This appeared to be slightly more pronounced at 18 nM Dox compared to 5 nM Dox. In sum, pulse-treatment of iPSCs with moderate doses of Dox is sufficient to impair mitochondrial function, and that this impairment is not reversed within the recovery period of 48 hours. As the same conditions do not appear to induce significant alterations in nuclear DDR or apoptosis, we conclude that these drug treatment conditions may selectively impair mitochondrial functionality. We propose that with these findings it is now possible to dissect known mitochondrial from nuclear effects supporting the view that mitochondria are a critical target of Dox.

**Figure 3.**
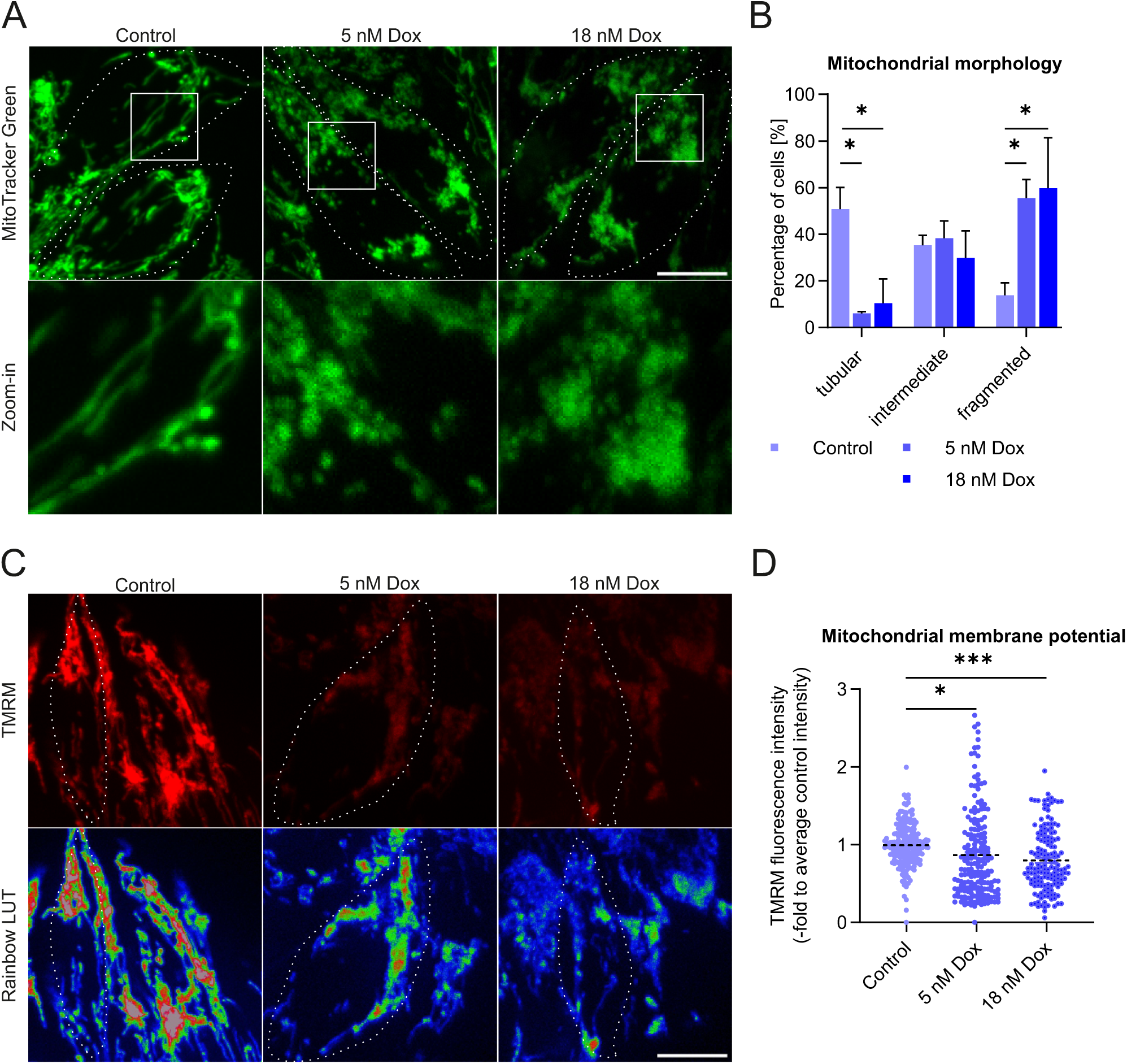
Doxorubicin pulse-treatment of iPSCs alters mitochondrial morphology and reduces the mitochondrial inner membrane potential. **A** Representative confocal image of MitoTracker™Green FM-stained mitochondria of control and pulse-treated iPSCs. Magnification: x63, scale bar: 10 μm. Cell surroundings are marked with dashed lines. **B** Quantification of mitochondrial morphology categorized in tubular, intermediate, or fragmented 48 hours after Dox treatment. Data show mean ± SEM (n=3; N=50 cells). 2-way ANOVA was performed. * p-value ≤ 0.05. **C** Representative confocal image of TMRM stained iPSCs (upper row). Intensity is shown in pseudo-color rainbow LUT from confocal images (lower row). Magnification: x63, scale bar: 10 µm. **D** Quantification of relative TMRM fluorescence intensity in Dox-treated iPSCs relative to the untreated control. Data represents the median plus whiskers (n=3; N=35-50 cells). For statistical analysis, one-way ANOVA with multiple comparisons was performed. * p-value ≤ 0.05. *** p-value ≤ 0.001.

### Pulse-treatment of iPSCs with doxorubicin impairs mitochondrial respiration and ATP formation

Since loss of mitochondrial membrane potential could result from impaired mitochondrial respiration, live cell respirometry was performed using a Seahorse Flux Analyzer. We observed a significant reduction in the overall oxygen consumption rate (OCR), maximal respiration (Max) and ATP production upon treatment with Dox (Fig. 4A, B). To further validate the Dox-mediated decrease in ATP production, steady-state levels of ATP were measured with the CellTiter-Glo® assay. With increasing Dox concentrations, ATP production significantly decreased reaching a reduction of about 20 % when 18 nM Dox were applied (Fig. 4C). Furthermore, we asked whether the suppression of mitochondrial respiration was linked to changes in mitochondrial ultrastructure. For this, transmission electron microscopy was performed and crista were categorized into four main morphologies: Tubes, septa, arches and branches. No significant changes in the cristae number per mitochondria after pulse-treatment with Dox were observable (Fig. 4D, E). Next, the influence of low Dox doses on the assembly of oxidative phosphorylation (OXPHOS) complexes was determined via performance of BN- and CN-PAGE analysis. Short pulse-treatment of iPSCs with Dox had no significant effect on the steady-state assembly of different OXPHOS complexes or the MICOS complex in iPSCs (Fig. 4F, G). However, it is noteworthy that iPSCs clearly harbor supercomplexes, high-molecular weight complexes composed of at least two different OXPHOS complexes (Fig. 4F, G), an observation that, to our knowledge, was not reported before. Measurement of in-gel activity of complex I and complex IV showed no significant effect of Dox on their activities. Overall, pulse-treatment showed a clear reduction in mitochondrial respiration, however, this was apparently not a result of altered mitochondrial ultrastructure or OXPHOS complex assembly or activity.

**Figure 4.**
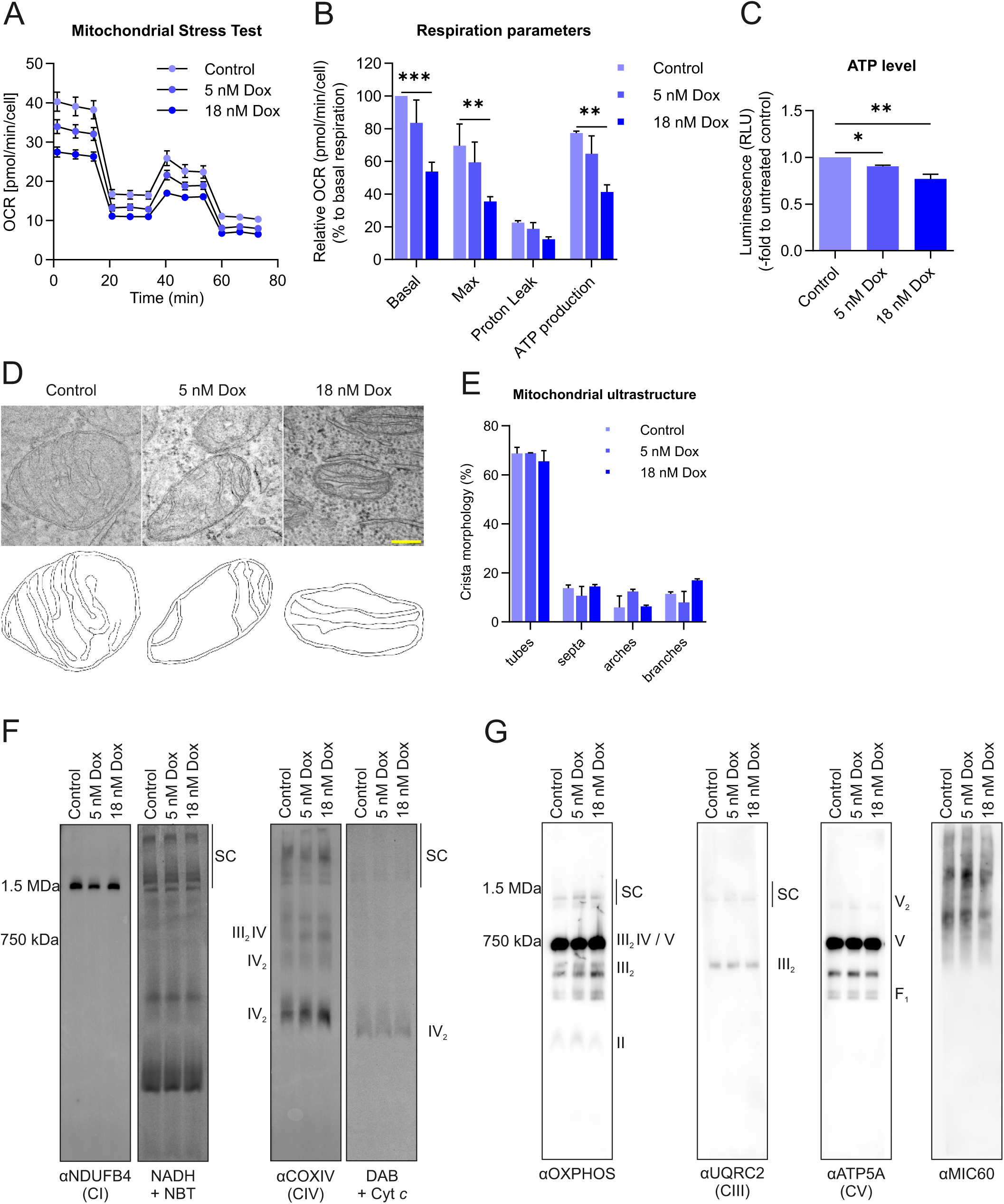
Mitochondrial ultrastructure and OXPHOS complexes are not affected by short pulse-treatment with doxorubicin. **A** Representative Seahorse run of one biological replicate. Shown is the oxygen consumption rates (pmol O_2_ / min) normalized to the cell number (by Hoechst staining). **B** Relative basal respiration, maximal respiration (after FCCP injection), proton leak and ATP production are shown for iPSCs after 48 hours of 2 hours pulse-treatment with Dox. Data was normalized to the untreated control and represents the mean ± SEM (n=3). For statistical analysis, *2-way* ANOVA with multiple comparisons was performed. ** p-value ≤ 0.01, *** p-value ≤ 0.001. **C** Quantification of relative ATP levels 48 hours after 2 hours pulse-treatment with Dox using CellTiter-Glo® 2.0 Cell Viability Assay. Data represent the mean ± SEM (n=3; N=4), normalized to respective control. *One-way* ANOVA with multiple comparisons was performed. * p-value ≤ 0.05, ** p-value ≤ 0.01. **D** Representative electron micrograph of mitochondria from iPSC upon 5 nM and 18 nM Dox treatment. Drawn surroundings of respective mitochondria are shown in the second row. Scale bar 0.1 µm. **E** Quantification of crista per mitochondria 48 h after Dox treatment. Percentage of crista morphology categorized in tubes, septa, arches and branches in iPSCs after Dox treatment. Data show mean ± SEM (n=2; N= 38-50 mitochondria). **F** Blue (left) and clear native (right) PAGE analysis of ETC complex I (CI) and IV (CIV) in control and Dox pulse-treated iPSCs. Shown are representative images of blue and clear native PAGEs (blue native PAGE: n=4; clear native PAGE: n=2). **G** Blue Native PAGE gels blotted and probed with indicated OXPHOS complex II (CII), III (CIII), V (CV) and Mic60 antibodies upon pulse-treatment with Dox in iPSCs. Representative images are shown (n=4).

### Pulse-treatment with doxorubicin modulates organization of the mitochondrial genome in iPSCs

Within mitochondria, several hundreds to thousands of mtDNA copies are organized as punctate-like nucleoid structures harboring ∼1.4 mtDNA molecules per nucleoid on average (Kukat et al., 2011). To determine the effect of Dox on mitochondrial nucleoid organization, mitochondrial nucleoid content and morphology was determined using confocal microscopy. Upon Dox treatment, the average number of mitochondrial nucleoids per cell was reduced with both concentrations of Dox compared to the control, even showing a slightly more pronounced effect when using 5 nM Dox (Fig. 5A, B). Next, we determined the average size of mitochondrial nucleoids. Pulse-treatment of iPSCs is accompanied with a moderate but significant reduction in nucleoid size of nucleoid size when 5 nM Dox were applied compared to the untreated control as well as to the 18 nM Dox treated condition (Fig. 5C). At 18 nM Dox there was no significant change of the average size compared to the control. To elucidate whether Dox-dependent modulation of mitochondrial nucleoids results in overall reduction in mitochondrial DNA content (mtDNA copy number) was measured by quantitative real-time PCR (Fig. 5D). Upon treatment of iPSCs with Dox, only a moderate, yet not significant, increase in mtDNA copy number was found. In summary, the administration of pulse-treatment with Dox in human iPSCs has been demonstrated to induce a modulation in the organization of mitochondrial DNA, characterized by a moderate change in the size of mitochondrial nucleoids, concurrent with a decrease in nucleoid number. This finding is consistent with the possibility that mitochondrial nucleoids may actually undergo remodeling in response to Dox-dependent inhibition of mtDNA replication.

**Figure 5.**
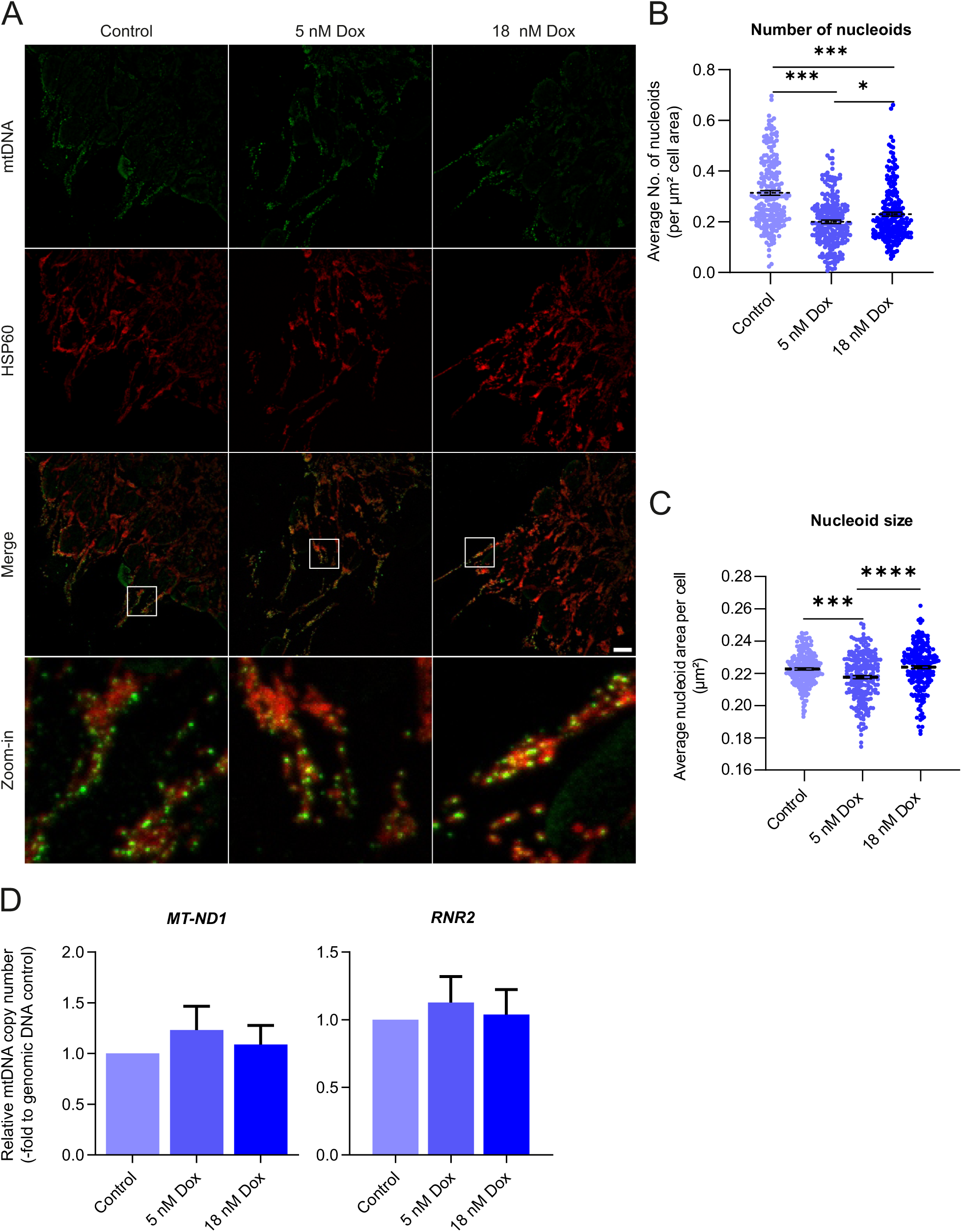
Pulse-treatment of iPSCs with the genotoxic anthracycline doxorubicin alters mtDNA organization. **A** Representative confocal image of mitochondrial nucleoids (α-DNA) in iPSC upon 5 nM and 18 nM Dox treatment. Magnification: x 63, scale bar: 10 μM. Zoomed regions are marked with a white square. **BC** Quantification of the average amount of mitochondrial nucleoids per µm² per cell (**B**) and nucleoid size in µm² (**C**) after pulse-treatment with Dox. Data show mean ± SEM (n=3; N=50 cells). *One-way* ANOVA was performed. * p-value ≤ 0.05,** p-value ≤ 0.01, ***p-value ≤ 0.0001. **D** Quantitative real-time PCR analysis of mitochondrial encoded genes mitochondrial encoded 16S rRNA (*RNR2*) and NADH-ubiquinone oxidoreductase chain 1 (*MTND1*) in pulse-treated iPSCs. Relative expression levels of three independent biological experiments are shown (n=3; N=3).

### Pulse-treatment of human iPSCs with doxorubicin results in major changes in gene expression modulating cellular homeostasis and metabolism

Since we observed that pulse-treatment of iPSCs with Dox impairs mitochondrial functions, morphology and mtDNA organization, we asked which transcriptional response on a genome-wide level is induced by this treatment and whether metabolic pathways linked to mitochondria are affected. For this, we performed whole genome transcriptome profiling of human iPSCs 48 hours after 2 hours pulse-treatment with Dox and the respective mock-treated iPSC control (Fig. 6A). Multidimensional scaling revealed high homogeneity within the respective replicates (Supplementary Figure 4). In total the expression levels of 16198 genes were obtained from the short read-NGS RNAseq dataset. Pairwise comparison between pulse-treated iPSCs (iPSC_Dox) and the respective control (iPSC_Mt) resulted in significant upregulation of 753 and downregulation of 519 genes (Supplementary Table 2). A GO-term enrichment analysis revealed no significant functions in the genes upregulated after Dox-treatment of iPSCs, whereas, for the significantly downregulated genes an enrichment of various biological processes (BP) was seen e.g. cellular response to interleukin-7, cytoplasmic translation, negative regulation of protein modification, and pyruvate metabolic process (Fig. 6B). Several ribosomal genes and eukaryotic initiation factors involved in cytoplasmic translation were significantly downregulated as well as Lin-28 Homolog A (*LIN28A*), a pivotal regulator of genes enhancing pluripotency in stem cells and modulating metabolism (Peng et al., 2011; J. Zhang et al., 2016). Additionally, genes involved in post-translational regulation, including the Ubiquitin Protein Ligase E3 Component N-Recognin 5 (*UBR5*) and Sirtuin 1 (*Sirt1*) were found to be downregulated. In addition, pulse-treatment of iPSCs was accompanied with suppression of SRY-Box Transcription Factor 4 (*SOX4*), a crucial transcription factor during embryogenesis (Fig. 6B). This suggests that even transient Dox exposure may disrupt cellular protein homeostasis and cell fate decisions. Importantly, within the group of downregulated genes we also observed numerous factors controlling major metabolic pathways such as hexokinase 1 (*HK1*), pyruvate dehydrogenase kinase 1 (*PDK1*), phosphoglucomutase 1 (*PGM1*), pyruvate dehydrogenase E1 subunit beta (*PDHB*), and hypoxia inducible factor 1 subunit alpha (*HIF1A*). These enzymes and factors clearly hint towards an alteration of glycolysis as well as downstream metabolic pathways occurring inside mitochondria upon Dox-treatment. In sum, pulse-treatment for only two hours with moderate doses of Dox is sufficient to cause major transcriptional changes accompanied with downregulation of pivotal genes regulating cellular homeostasis and energy metabolism in iPSCs.

**Figure 6.**
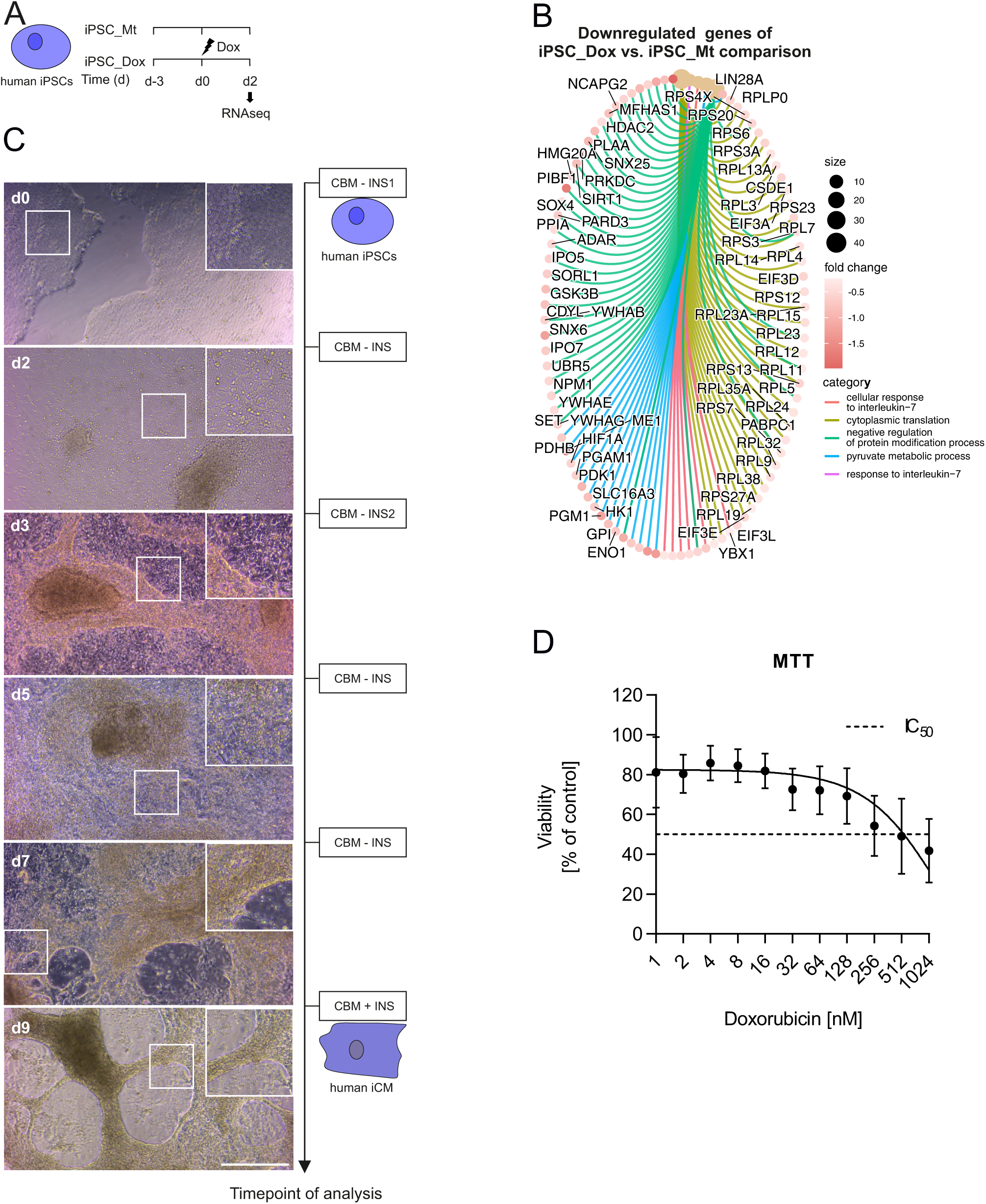
GO-term enrichments of differentially regulated genes in induced-pluripotent stem cells influenced upon pulse-treatment with doxorubicin. **A** Schematic illustration of the treatment scheme in iPSCs prior to short-read-NGS RNAseq analysis at day 2 (d2). Timepoint of 2 hours pulse-treatment with 18 nM doxorubicin (Dox) is indicated by a lightning symbol. **B** Gene concept network of GO-term enrichments and associated downregulated genes of pairwise comparison between Dox-treated iPSCs and the respective control. Number of genes, logFC-value, and GO-terms are listed in the respective legend. **C** Scheme of iPSC differentiation into iPSC-derived cardiomyocytes (iCMs) following a chemically defined protocol. At defined days (d), cardiac basal media (CBM) was supplemented without B-27 minus insulin (-INS) or with insulin (+INS). In addition, growth factors such as CHIR99021 (CBM-INS1) or IWP4 (CBM-INS2) were added at specific days of the differentiation protocol. Days with negative numbers represent the time of culturing iPSCs until initiation of cardiac differentiation. On day 7 differentiation of iPSCs into iPSC-derived cardiomyocytes (iCMs) is completed. Representative images of iCMs acquired with Leica DM IL LED Fluo Cellfactory (Leica). Days (d) of cardiac differentiation are indicated. Magnification: 20x, scale bar: 300 µm. **D** Determination of sensitivity of iCMs against Dox. iCMs were pulse-treated for 2 hours with increasing doses of doxorubicin (Dox) and viability was determined using an MTT assay. The mean ± SEM of three independent biological replicates (n=3; N=4-8) is shown.

### iPSCs-derived cardiomyocytes (iCMs) show a strongly increased resistance against doxorubicin compared to iPSCs

Based on the observed Dox sensitivity of iPSCs, we next asked whether iPSCs can be differentiated into iPSC-derived cardiomyocytes (iCMs) and whether the sensitivity of iCMs towards Dox differs from their undifferentiated counterparts. For this, a previously established differentiation protocol was applied (Fig. 6C) and the generated iCMs were characterized. Consistent with a successful cardiac differentiation we observed a cell morphology resembling cardiomyocytes at day 9 of the differentiation protocol (Fig. 6C). Generated iCMs further showed spontaneous contractions starting from day 7 of the cardiac differentiation protocol (Supplementary movies 1-4). Cardiac differentiation was corroborated by quantitative real-time PCR showing enhanced expression of cardiomyocyte markers Troponin T2 (*TNNT2*), NK2 Homeobox 5 (*NKX2-5*), Myosin Heavy Chain 6 (*Myh6*) and GATA Binding Protein 4 (*GATA4*) as well as reduced expression of pluripotency markers POU Class 5 Homeobox 1 (*POU5F1*), Nanog Homeobox (*NANOG*) and SRY-Box Transcription Factor 2 (*SOX2*) (Supplementary Figure 3). To determine the sensitivity of iCMs against doxorubicin in a comparable manner, we applied a similar 2-hour pulse-treatment scheme as in iPSCs and measured cell viability after 48 h using the MTT assay. Here, iCMs showed an IC_50-_value of ∼512 nM (Fig. 6D) which is much higher than the IC_50_ value of ∼35 nM for iPSCs. Consistent to the treatment scheme applied in iPSCs, in all subsequent experiments iCMs were treated with a Dox dose equivalent to IC_30_-value, namely 144 nM. In sum, iCMs are considerably more resistant against Dox compared to their undifferentiated iPSCs counterparts.

### The applied mild pulse-treatment scheme for Dox in iPSCs does not impair differentiation into iPSC-derived cardiomyocytes (iCMs)

Next, we asked whether pulse-treatment of iPSCs with moderate doses of doxorubicin affects their differentiation into iCMs, whether this has persistent transcriptional consequences after cells have differentiated into cardiomyocytes, and whether there are functional consequences. In other words, are iCMs that were already treated in iPSC stage transcriptionally preconditioned and does this impose an advantage or disadvantage compared to mock-treated cells when coping with similar types of stress? To address this, the following four different treatment schemes were applied (Fig. 7A) and iCMs were analyzed together with Dox treated and mock-treated iPSCs (Fig. 6AB) by whole genome transcriptome profiling. For example, iPSCs were only pulse-treated with the corresponding IC_30_ dose (18 nM) before initiation of cardiac differentiation and analyzed at the iCM stage (iCM_Dox_Mt). To determine the influence of moderate pulse-treatment of Dox on terminally differentiated cardiomyocytes, iCMs were treated with the respective IC_30-_dose of 144 nM for 2 hours on day 7 of cardiac differentiation followed by 48 hours recovery (iCM_Mt_Dox). To investigate whether early pulse-treatment of iPSCs alters the response towards Dox treatment at a later stage of cardiac differentiation Dox-treated iPSCs were exposed to a second dose of Dox at the iCM stage (iCM_Dox_Dox). Here, each of the three treatment schemes (iCM_Dox_Mt, iCM_Mt_Dox, iCM_Dox_Dox) were compared to a mock-treated control (iCM_Mt_Mt). Whole genome transcriptome profiling revealed numerous significantly differentially expressed genes (Supplementary Table 2) and the results and implications are discussed in a systematic manner in the following sections.

**Figure 7.**
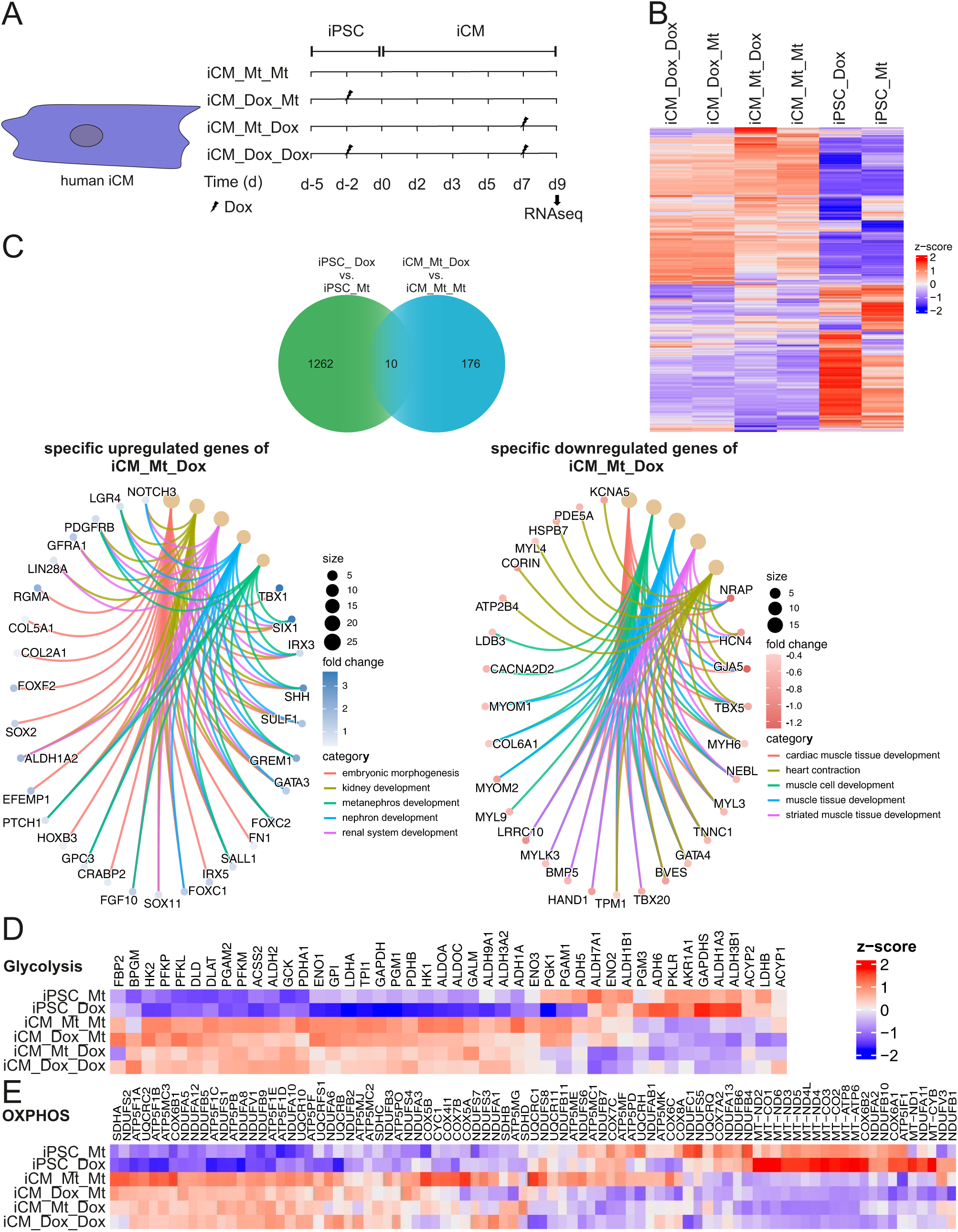
Whole-genome transcriptome analysis reveals differences in the response of iPSCs and iCMs towards doxorubicin treatment. **A** Overview of treatment schemes in iCMs prior to whole genome-RNAseq analysis. Following samples were measured: Untreated iCMs (iCM_Mt_Mt), in iPSC stage treated iCMs (iCM_Dox_Mt), iCM treated on day 7 (iCM_Mt_Dox) and iCMs treated in iPSC stage and on day 7 of cardiac differentiation (iCM_Dox_Dox). Timepoint of pulse treatment with doxorubicin (Dox) is marked with a lightning symbol. **B** Heatmap of all 16198 genes detected in the indicated six treatment conditions. The average z-score is given of all four technical replicates in all conditions scaled per gene. **C** Venn diagram of pairwise comparisons between Dox-treated iPSCs and iCMs pulse-treated with Dox on day 7 of cardiac differentiation compared to the respective control. Gene concept network of GO-term enriched upregulated (top panel) and downregulated (bottom panel) genes of pairwise comparison between Dox-treated iPSCs (iPSC_Dox) and iCMs (iCM_Mt_Dox). Number of genes, logFC-value, and GO-terms are indicated. **DE** Heatmap of targeted analysis of genes involved in glycolysis and gluconeogenesis (**D**) and oxidative phosphorylation (OXPHOS, **E**). Average z-score for all four technical replicates of each condition are shown. Coloring for z-scores as indicated.

The first observation is that differentiation into iCMs is not impaired independent of the timepoint of Dox pulse-treatment as cellular morphology and beating of iCMs appears normal (Supplementary Movies 1 to 4). Also, the observed expression levels of cardiac differentiation and pluripotency markers by quantitative real-time PCR corroborate this finding (Supplementary Figure 3). Moreover, multidimensional scaling of the transcriptome data showed a clear trend of separation between the six groups with a high homogeneity within the replicates (Supplementary Figure 4). Overall, data for 16198 genes were received from the RNA-seq data set derived from iPSCs and iCMs. By performing hierarchical clustering, we found strong differences in gene clusters between iPSCs and their differentiated counterparts reflecting the major transcriptional changes induced by the differentiation process itself (Fig. 7B; Supplementary Figure 6). Thus, independent of the timepoint of Dox treatment, the expression of typical cardiac differentiation markers was also seen on a whole transcriptome level confirming that differentiation of iPSCs into cardiomyocytes proceeds despite prior or concurrent Dox exposure.

### The transcriptional response to pulse-treatment with doxorubicin reveals major differences between iPSCs and iCMs

Given that both untreated and Dox-treated iPSCs can differentiate into iCMs, we next asked how iCMs are affected by a pulse-treatment with doxorubicin on a global transcriptome level and whether the response is different to the Dox effect in iPSCs described earlier. For this, pairwise comparisons between Dox-treated iPSCs and iCMs on day 7 of cardiac differentiation to their respective control was performed (iPSC_Dox vs. iPSC_Mt; iCM_Mt_Dox vs. iCM_Mt_Mt) to determine significantly differentially regulated genes upon Dox treatment (Fig. 7C; Supplementary Table 2). A subsequent pairwise comparison between the resulting gene sets revealed a total of 842 significantly upregulated and 606 downregulated genes (Fig. 7C). Interestingly, only 10 significantly differentially regulated genes were shared between both conditions, whereas most changes were specific to the cell type treated. Importantly, iPSCs showed a higher number of specific genes (1262) significantly affected by moderate pulse-treatment with Dox as compared to their differentiated counterparts (176) supporting the hypothesis that Dox has a major impact on gene expression specifically in iPSCs. GO-term enrichment of biological processes (BP) analysis of genes solely affected in iPSCs (1262) resulted in the enrichment of the same GO-terms as described previously for Dox-treated iPSCs (Fig. 6B). In contrast to iPSCs, the number of specific and significantly affected genes in iCMs was much lower, with 89 upregulated genes and 87 downregulated genes. An analysis of those significantly upregulated genes in Dox-treated iCMs revealed GO-term enrichment of biological processes (BP) linked to embryonic morphogenesis and kidney development (Fig. 7C). Among these GO-term functional enrichments associated genes such as T-Box transcription factor 1 (*TBX1*), SIX homeobox 1 (*SIX1*), sonic hedgehog signaling molecule (*SHH*), and repulsive guidance molecule BMP co-receptor A (*RGMA*) have shown to be highly upregulated indicating Dox-induced upregulation of processes associated with embryogenesis. As shown by GO-term enrichment analysis, pulse-treatment with Dox induces the repression of genes associated with cardiac muscle development, heart contraction, muscle cell development, muscle tissue development, and striated muscle tissue development (Fig. 7C). Here, gap junction protein alpha 5 (*GJA5*), nebulin related anchoring protein (*NRAP)*; and hyperpolarization activated cyclic nucleotide gated potassium channel 4 (*HCN4*) were suppressed upon pulse-treatment with Dox. Suppression of these genes are linked to impaired formation of electrochemical gradients and cellular connectivity necessary for cardiac function. Taken together, iPSCs and iCMs show major differences in the transient transcriptional response upon Dox pulse-treatment. Pulse-treatment of iCMs with Dox was not accompanied with major significant GO-terms associated with metabolism as seen in iPSCs pointing to distinct effects of DOX in iCMs versus iPSCs. Since we could show that pulse treatment of iPSCs is accompanied with downregulation of metabolic function (Fig. 4A-C), we questioned whether this effect is reflected in our transcriptomic data. A targeted analysis using a defined gene set for glycolysis and gluconeogenesis from Harmonizome 3.0 (Diamant et al., 2025; Rouillard et al., 2016) revealed a Dox-induced reciprocal transcriptional response of essential genes of glycolysis in iPSCs versus iCMs (*HK1, GAPDH, ALDOA, ALDOC, LDHA*, *PGK1*) independent of the timepoint of treatment (Fig. 7D, Supplementary Figure 6). Similar reciprocal effects were observable when analyzing a defined set of genes for oxidative phosphorylation (OXPHOS) from Harmonizome3.0 (Fig. 7E, Supplementary Figure 6). It is noteworthy that iPSCs showed a very robust upregulation of all 13 mitochondrial encoded genes (*MT-ATP6, MT-ATP8, MT-CO1, MT-CO2, MT-CO3, MT-CYB, MT-ND1, MT-ND2, MT-ND3, MT-ND4, MT-ND4L, MT-ND5, MT-ND6*) which was clearly not seen in iCMs.

In sum, moderate single pulse-treatment of iPSCs and iCMs revealed major differences in the transcriptional response for the two differentiation states. While the transcriptional response in iCMs clearly modulated gene expression mediating cardiac function, the response of iPSCs revealed many genes altering major metabolic processes such as glycolysis and OXPHOS. In iCMs, the latter effect was not evident or appeared even reciprocal. Thus, even modest pulse-treatment in iPSCs is accompanied with upregulation of mtDNA-encoded genes supporting the hypothesis that this could represent a compensatory mechanism to balance mitochondrial dysfunction upon treatment with Dox.

### A single early pulse-treatment with Dox in iPSCs causes transcriptional reprogramming in resulting iCMs associated with impaired cardiac muscle regeneration

Next, we asked whether Dox pulse-treatment in the iPSC stage causes lasting changes on the transcriptome level in the resulting iCMs. To analyze this, we performed a pairwise comparison between iCMs treated prior to their differentiation with Dox in their iPSC-stage and their respective control (iCM_Dox_Mt vs. iCM_Mt_Mt). Whole genome transcriptome analysis revealed upregulation of 233 and downregulation of 193 genes (Supplementary Table 2). GO-term enrichment showed a significant upregulation of genes linked to biological processes (BP) such as angiogenesis, blood vessel morphogenesis, positive regulation of cell migration, and positive regulation of cell motility (Fig. 8A). Among these terms, several genes regulating and promoting collagen production have been detected consistent with a pro-fibrotic shift in gene expression. Conversely, downregulated genes were enriched in GO-terms related to cardiac muscle tissue development, heart development, and muscle tissue development (Fig. 8B).

**Figure 8.**
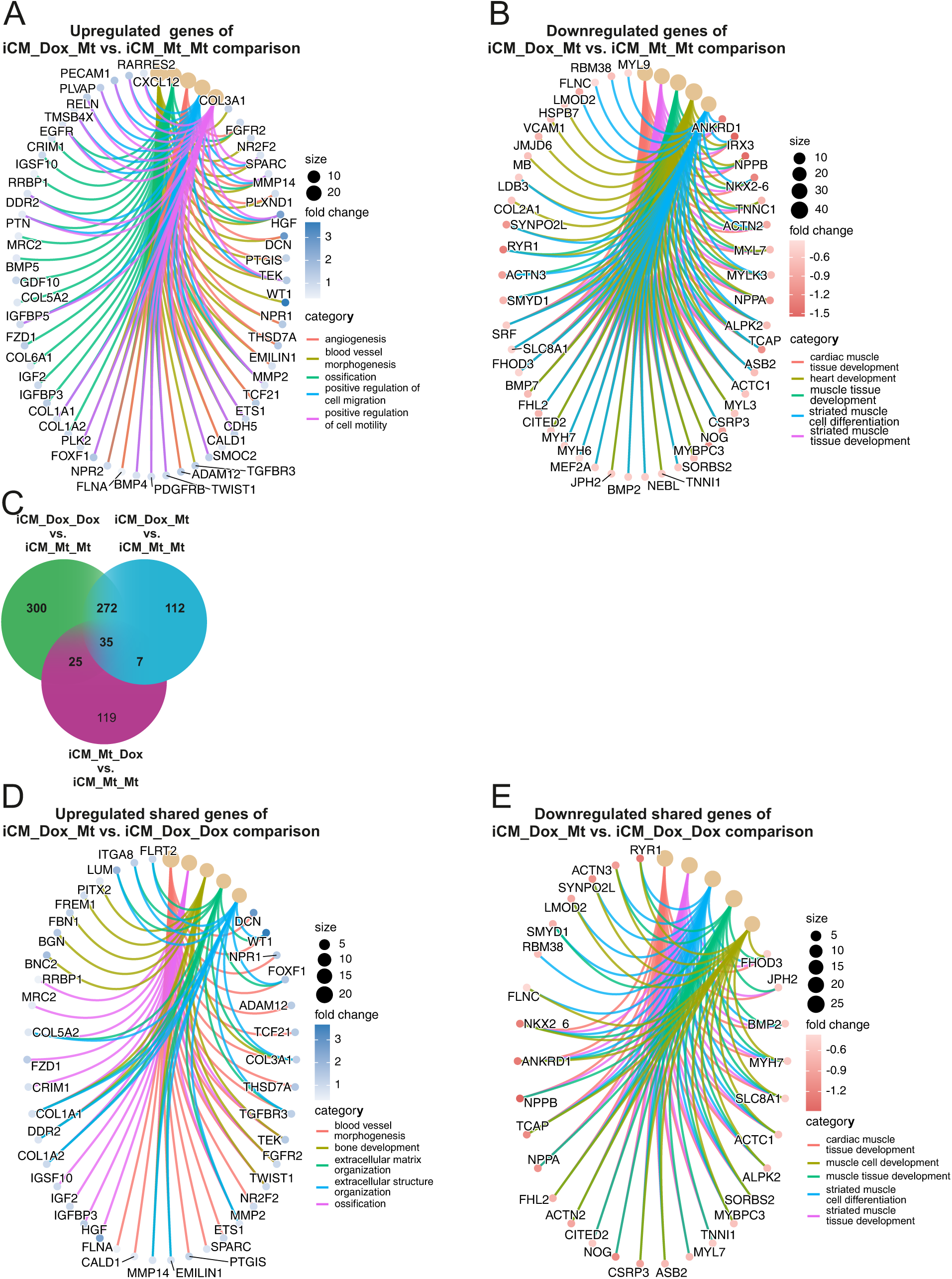
Early doxorubicin treatment of iPSCs is accompanied with whole genome transcriptional reprogramming in iCMs. **AB** Gene concept network of GO-term enrichments of differentially upregulated (**A**) and downregulated (**B**) genes of iCMs (iCM_Dox_Mt) compared to mock-treated control (iCM_Mt_Mt). GO-terms of associated genes, number of genes, and logFCs are indicated. **C** Venn diagram of significantly differentially regulated genes resulting from indicated pairwise comparisons in which mock-treated iCMs (iCM_Mt_Mt) were always used as control. **DE** Gene concept network of GO-term enrichments and associated shared upregulated (**D**) and downregulated (**E**) genes when early treatment of iPSCs occurred (iCM_Dox_Mt and iCM_Dox_Dox). Number of genes, GO-terms, and logFCs are shown.

To gain further information on the importance of the timepoint when Dox treatment occurred, we performed various pairwise comparisons of all treatment conditions in iCMs (iCM_Dox_Mt, iCM_Mt_Dox, and iCM_Dox_Dox all compared to the non-treated control iCM_Mt_Mt; Fig. 8C).

Initially, we focused to elucidate the influence of Dox treatment in the iPSC stage on the transcriptional response in iCMs. For that, we performed pairwise comparison of treatment conditions sharing an early Dox treatment in iPSC stage (iCM_Dox_Mt and iCM_Dox_Dox both compared to the non-treated control iCM_Mt_Mt) (Fig. 8C). Interestingly, 272 differentially expressed genes were specifically shared, of which 139 genes were upregulated and 133 were downregulated (Fig. 8C, Supplementary Table 3). GO-term enrichment of biological processes (BP) revealed that upon early pulse-treatment with Dox, again genes involved in blood vessel morphogenesis, ossification, extracellular matrix organization, and extracellular structure organization were robustly upregulated (Fig. 8D; Supplementary Figure 6), namely independent of a Dox-treatment applied in the iCM stage, emphasizing that this transcriptional response is caused before differentiation was initiated. Key genes included Decorin (*DCN*), WT1 Transcription Factor (*WT1*), Filamin A (*FLNA*), and Basonuclin Zinc Finger Protein 2 (*BNC2*), which all have been described to play pivotal roles in regulation of fibrosis and cardiac regeneration. However, treatment of cells in iPSC stage is accompanied with strong downregulation of genes associated with cardiac muscle tissue development, striated muscle cell differentiation, muscle tissue development, and muscle cell development (Fig. 8E; Supplementary Figure 6). Among these GO-term functional enrichments, NK2 Homeobox 6 (*NKX2-6*), Ryanodine Receptor 1 (*RYR1*), Ankyrin Repeat Domain 1 (*ANKRD1*), and Actinin Alpha 3 (*ACTN3*) have shown to be the most suppressed genes upon Dox treatment before cardiac differentiation. All these mentioned genes are essential regulators of cardiac function. We conclude that treatment with a moderate and transient dose of Dox at an iPSC stage prior to cardiac differentiation is sufficient to cause a major transcriptional reprogramming persistent in differentiated cells altering blood vessel formation and promoting fibrosis, while expression of genes required for cardiac muscle regeneration are repressed. This would be in line with a reduced resulting regenerative capacity when iPSCs have been exposed to even low and transient doses of Dox once before the differentiation or regeneration process has been initiated.

### Pulse-treatment of iCMs results in transcriptional adaptations promoting apoptosis

Based on the major impacts seen on the expression profile when cells were pulse-treated prior to cardiac differentiation, we asked how a ‘late’ Dox pulse-treatment at the iCM stage, as opposed to the iPSC stage, influences the transcriptional response in terminally differentiated iCMs. For this, we focused on shared genes that are significantly differentially regulated when iCMs were treated late (iCM_Mt_Dox and iCM_Dox_Dox, both compared to iCM_Mt_Mt (Fig. 8C). The number of shared genes differentially regulated upon late treatment was much lower compared to the shared genes upon early treatment (Fig. 8C, Supplementary Table 3). Only 25 genes were specifically shared, of which 12 genes were upregulated and 13 genes were downregulated (Fig. 8C, Supplementary Table 3). GO-term enrichment of biological processes (BP) showed an enrichment in genes associated with positive regulation of programmed cell death, positive regulation of apoptotic process and apoptotic signaling pathway (Fig. 9A). Among these functional enrichments Fas cell surface death receptor (*FAS*), cyclin dependent kinase Inhibitor 1A (*CDKN1A*), and BCL2 associated X (*BAX*) were among the top candidates of upregulated genes. All of these genes have a regulative role in cell survival or cell cycle progression. Overall, exposure of terminally differentiated iCMs to a modest dose of Dox is accompanied with a transcriptional upregulation of apoptotic pathways consistent with a direct toxicity of Dox to iCMs promoting apoptosis. The fact that we do not see this apoptotic signature in iPSCs treated with Dox is consistent with our results described earlier showing no induction of extensive DDR or apoptosis.

**Figure 9.**
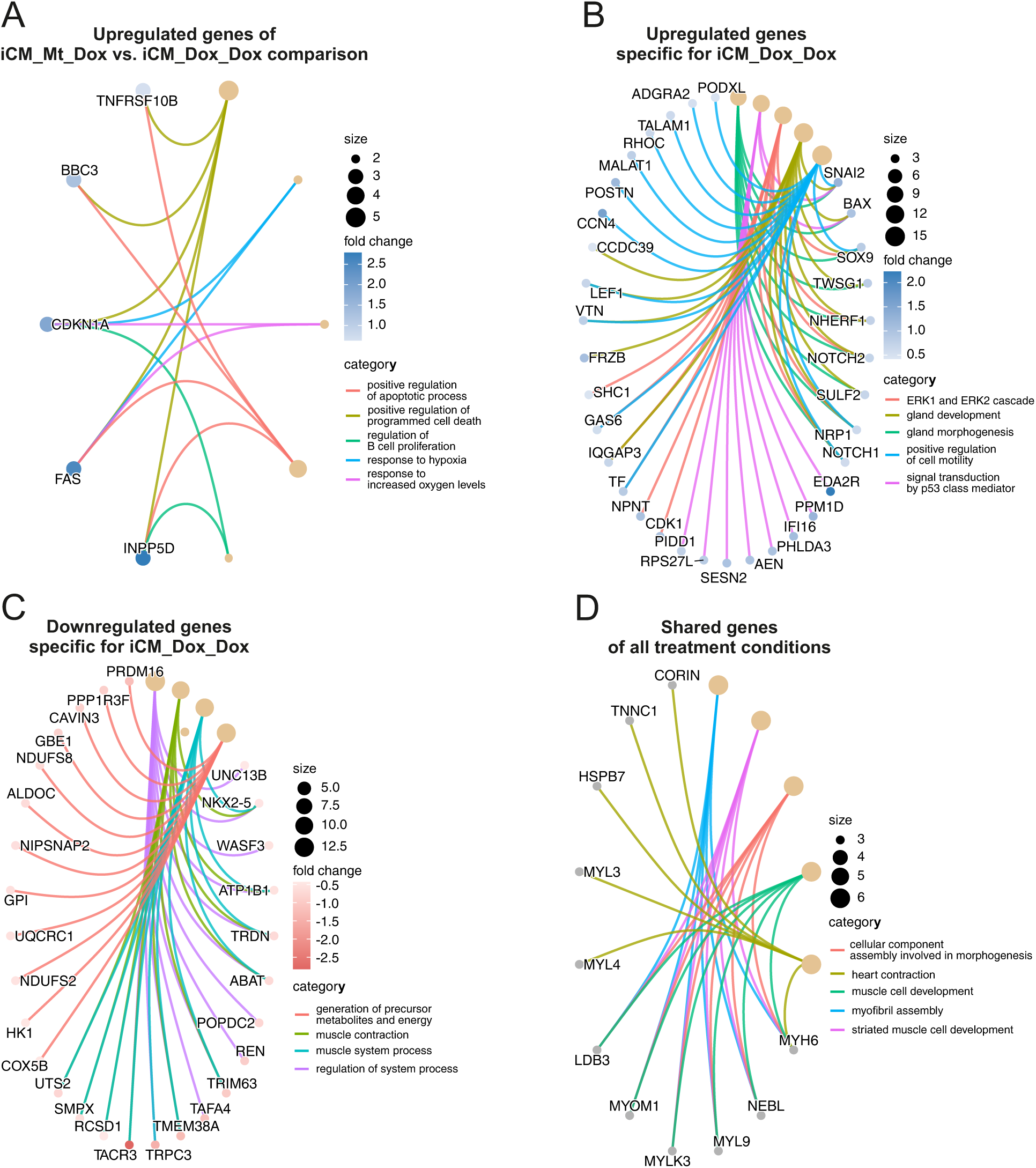
Doxorubicin applied prior to cardiac differentiation influences transcriptional response of iCMs towards doxorubicin applied later. **A** Gene concept network of GO-term enrichments of differentially upregulated genes exclusively shared by late-treated iCMs (iCM_Mt_Dox and iCM_Dox_Dox, all compared to iCM_Mt_Mt). GO-terms of associated genes, number of genes, and logFCs are indicated. **BC** Gene concept network of GO-term enrichments of differentially upregulated (**B**) and downregulated (**C**) genes unique for double-treated iCMs (iCM_Dox_Dox). GO-terms of associated genes, number of genes, and logFCs are indicated. **D** Gene concept network of GO-term enrichments and associated downregulated genes within the exclusive intersection of all treatment conditions in iCMs (iCM_Dox_Mt vs. iCM_Mt_Dox vs. iCM_Dox_Dox). Number of genes and GO-terms are indicated.

### The transcriptional response of iCMs to doxorubicin depends on an early pulse-treatment of Dox at the iPSC stage altering energy metabolism and muscle contraction pathways

Since iCMs that have been exposed to Dox before cardiac differentiation showed a strong transcriptional response, we questioned whether this early pulse-treatment alters the stress response against Dox when iCMs were exposed for a second time. Thus, we introduced a double-treatment condition including both treatment timepoints (Fig. 7A). We find 300 genes that are only differentially regulated upon double-treatment with Dox (iCM_Dox_Dox), of which 164 genes were upregulated and 136 genes were downregulated (Fig. 8C, Supplementary Table 3). In contrast, iCMs that only were exposed late to Dox (iCM_Mt_Dox) showed 119 different genes specifically affected emphasizing that the transcriptional response to a late Dox treatment is highly influenced by a type of preconditioning prior to commitment to cardiac differentiation. This is also corroborated by the fact that only 25 genes showed a specific common response upon late treatment. GO-term analysis of significantly differentially expressed genes upon a second Dox treatment solely found in pre-conditioned iCMs (iCM_Dox_Dox) showed significantly upregulated genes enriched for biological processes (BP) such as ERK1 and ERK2 cascade, positive regulation of cell motility, signal transduction by p53 class mediator, and gland development (Fig. 9B). Representative genes included cyclin dependent kinase 1 (*CDK1*), Wild-Type P53-induced phosphatase 1 (*PPM1D*), snail family transcriptional repressor 2 (*SNAI2*), and SRY-Box transcription factor 9 (*SOX9*), which have critical regulatory roles in cell fate decisions and transcription. However, pre-conditioning of iCMs already in iPSC stage was accompanied with downregulation of genes associated with energy metabolism, muscle system processes and muscle contraction (Fig. 9C). Within these GO-term functions, genes such as hexokinase 1 (*HK1*), several respiratory chain complex subunits (*COX5B*, *UQCRC1*, *NDUFS2*, *NDUFS8*), NK2 homeobox 5 (*NKX2-5*), ATPase Na+/K+ transporting subunit beta 1 (*ATP1B1*), and triadin (*TRDN*) were downregulated upon pulse-treatment with Dox. Dysregulation of these genes indicates a possible disturbance in energy metabolism and regulation of electrochemical gradients necessary for cardiac function. Overall, we conclude that early doxorubicin treatment at the iPSC stage causes not only a major reprogramming of iCMs themselves but also alters their transcriptional response to a second exposure to Dox markedly negatively impacting pathways linked to energy metabolism and muscle contraction.

### Pulse-treatment of Dox is generally accompanied with dysregulation of cardiac muscle formation in iCMs independent of the time when Dox was applied

To determine whether Dox exerts a common transcriptional response which might be shared by independent of the time or differentiation status of exposure, we identified DEGs commonly regulated over all treatment conditions (iCM_Dox_Mt, iCM_Mt_Dox, iCM_Dox_Dox) each compared to the mock-control (iCM_Mt_Mt). In total for all pulse-treatment schemes of Dox, 870 significantly differentially regulated genes were found (Fig 8C). Interestingly, only 35 genes were shared between all the treatment conditions, indicating a common Dox-specific effect on gene expression independent of the timepoint or differentiation status of treatment (Supplementary Table 3). Within the intersection of 35 selectively shared genes, some genes were antagonistically regulated as shown in Supplementary Figure 5. Enrichment analysis of GO-terms of biological processes (BP) showed no significant term enriched in the number of significantly upregulated genes. In contrast, GO-term enrichment analysis of downregulated genes revealed enrichment of genes associated with the GO-terms myofibril assembly, striated muscle cell development, heart contraction, and heart process (Fig. 9D). Among these enriched functions, Myosin Heavy Chain 6 (*MYH6*), Nebulette (*NEBL*) and Myosin Light Chain 9 (*MYL9*) were downregulated upon pulse treatment indicating a time-independent adverse effect of Dox treatment even upon modest and short exposure. In sum, pulse-treatment(s) with modest doses of Dox induce a convergent transcriptional downregulation of essential cardiac muscle genes in iCMs indicating an impairment of the contractile function of iCMs independently of the Dox exposure timing.

### Pulse-treatment of iPSCs with doxorubicin prior to differentiation results in functional changes in beating iCMs

To evaluate whether the transcriptional changes showing impaired cardiac function translate into functional impairments, we analyzed the spontaneous beating frequency of all four Dox treatment groups based on live-cell videos of iCMs (Fig. 7A) using the software Cardiovision. Overall, quantitative analysis revealed a significant reduction in the beating frequency in those iCMs that where ‘early’ pulse-treated with Dox at the iPSC stage and was most prominent in double-treated cells (Fig. 10A). Thus, preconditioning of iCMs appears to have a more profound functional effect than treating terminally differentiated iCMs with Dox that did not experience a Dox exposure at the iPSC stage. We next asked whether this early encounter with Dox might be accompanied with a higher sensitivity of iCMs against Dox. For this, cell viability with increasing concentrations of Dox was determined in cells preconditioned at the iPSC stage or for sham-treated cells using the SRB assay (Fig. 10B). Pulse-treatment of cells prior to cardiac differentiation was accompanied with an apparent moderately higher sensitivity against further Dox pulse-treatments at all Dox concentrations as compared to the mock-treated control, yet these differences were not significant.

**Figure 10.**
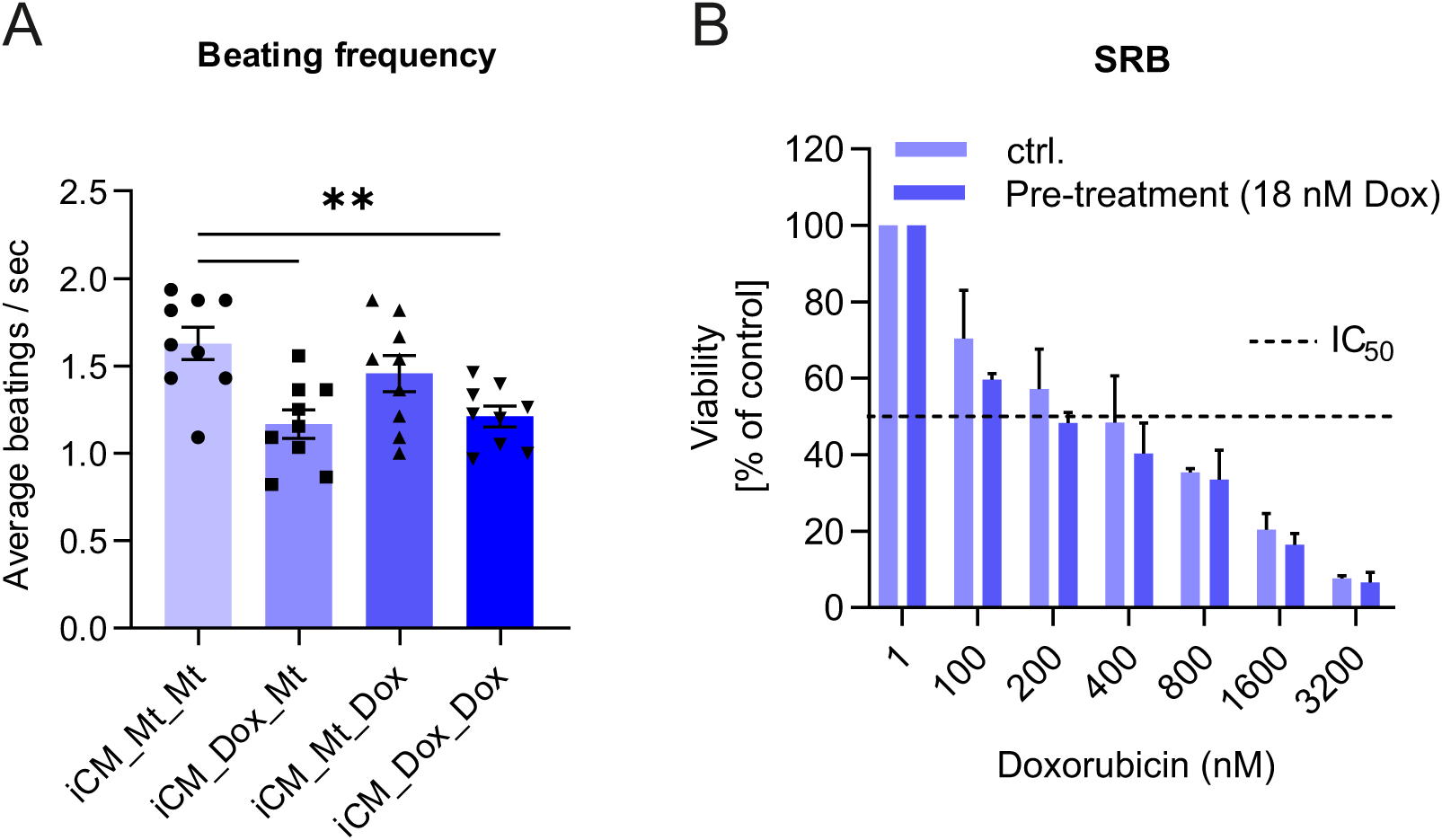
Repeated short-pulse treatment of cells with Dox is associated with impairment of cardiac beating parameters. **A** Average beating frequency per second measured in iCMs short-pulse treated with Dox at different stages of cardiac differentiation. Analysis of iCM contractility parameter in untreated iCMs (iCM_Mt_Mt), iCMs treated in iPSC stage (iCM_Dox_Mt), iCMs treated on day7 of cardiac differentiation (iCM_Mt_Dox) and in double-treated iCMs (iCM_Dox_Dox). iCMs were treated for 2 hours with Dox at defined stages of cardiac differentiation as shown in Fig. 9C. On day 9 of cardiac differentiation protocol, live-cell videos of about 20 seconds length with 60 frames per second of all conditions were acquired using Leica DM IL LED Fluo Cellfactory microscope. Analysis of the videos was conducted by using an automated analyzing system as described in the method section. Data shown represents the mean of three independent experiments ± SEM (n=3, N=3). The dashed line marks the IC_50_-value. One-way ANOVA with multiple comparisons was performed. ** p-value ≤ 0.01 **B** Measurement of diverse Dox concentrations on the viability of iCMs pre-treated in iPSC stage prior to cardiac differentiation and sham-treated cells. Data points shown represent the mean ± SEM of three independent experiments (n=3; N=8).

In accordance with our transcriptome findings, even short pulse-treatment with modest dose of Dox enhances dysregulation of cardiac contractility parameters independent of the timepoint of treatment. However, the strongest functional effect was observed in Dox double-treated cells indicating that Dox treatment at the iPSC stage negatively impacts a second encounter with Dox. We propose that preconditioned cells are somehow sensitized and that this is linked, at least in part, to mitochondrial dysfunction as well as to the major transcriptional reprogramming described here promoting fibrosis on the one hand side and impairing cardiac contraction function on the other side.

## Discussion

Dox is a widely-used and effective chemotherapeutic drug, but its clinical application is limited by its cardiotoxicity. Understanding the underlying pathomechanisms of this genotoxic drug is essential to develop strategies preventing heart failure and/or promoting heart regenerative processes. Here we have established and characterized a model system comprising of human iPSCs and iPSC-derived cardiomyocytes allowing us to study cellular and transcriptional consequences of moderate Dox pulse-treatment regimes. In particular, we aimed to understand the consequences of moderate Dox treatment conditions on mitochondrial and metabolic functions, how this depends on cardiac differentiation, and the impact of pre-conditioning at an iPSC sage on the susceptibility of cells differentiated from this. In contrast to many earlier studies, we applied a treatment scheme including rather mild and short exposures taking into account the low Dox concentrations reached rapidly in the plasma of treated patients receiving Dox as fast infusion (Gewirtz, 1999). Our study gives the following novel insights. First, human iPSCs are highly sensitive, much more than iCMs, to Dox. Exposure to low doses of Dox for only 2 hours was sufficient to reduce the viability of iPSCs significantly whereas iCMs were much more resistant. Second, this mild treatment primarily led to multiple defects in mitochondrial functionality including increased mitochondrial fragmentation, enhanced OPA1 processing, reduced inner membrane potential and altered mtDNA genome organization. The latter is consistent with the known genotoxic role of DOX by inhibiting mitochondrial DNA replication via intercalation as well as inhibition of the mitochondrial DNA-Topoisomerase IIβ. Importantly, it should be emphasized that these mild conditions did not result in activating a DDR or cell death, at least to a detectable manner, allowing to differentiate between toxic effects of DOX on mitochondrial DNA versus nuclear DNA. Third, upon analyzing the genome-wide transcriptional response to Dox at different time-points of differentiation we observed that iPSCs show an entirely different response to Dox than iCMs and much more genes were differentially expressed. Forth, another striking finding in our view is that iCMs treated at the iPSC stage, which we term preconditioning, have a very distinct transcriptional expression profile compared to iCMs that are not preconditioned. Interestingly, exactly those pathways that are linked to cardiac regeneration are reduced and pathways that promote fibrosis are increased in cells that have been exposed to Dox before differentiation. Also, metabolic pathways were differentially affected. Overall, this demonstrates for the first time that pluripotent, non-differentiated cells are genetically reprogrammed when exposed to a genotoxic substance in a way that does not alter differentiation *per se* but does alter the capacity to regenerate cardiac muscle. The observation that stem-cells exhibit high sensitivity against Dox exposure, albeit DDR is not induced, strongly support that primarily mitochondrial impairments lead to cellular dysfunction and suggests that mitochondria are one prime target for Dox, even at very low concentrations and in particular for pluripotent stem cells.

Here, the application of mild and transient exposures of a genotoxic compound revealed a number of unexpected insights compared to earlier studies using rather high doses for longer time periods. For example, a high susceptibility of stem cells to undergo programmed cell death was reported and has been proposed to ensure the maintenance of the stem cell pool by elimination of genetically defect cells (Fan et al., 2011). Moreover, stem cells were proposed to rely strongly on complex DNA repair machineries to encounter possible DNA damage (Rocha et al., 2013). We do not question these findings; however, it appears that this cannot be generalized for milder treatment conditions. At the conditions applied here, an induction of apoptosis upon treatment with Dox was not observed. Further investigation of DNA damage repair (DDR) of iPSCs in our setting revealed no detectable DDR induction or signs of DNA damage. Thus, the high sensitivity of iPSCs has to be based on other factors, yet not on induction of persistent DNA damage or insufficient DNA repair.

We clearly demonstrate, that mitochondria are one major target of Dox-induces and propose that they at least contribute to explain our findings. Besides the genomic DNA in the nucleus, mitochondria also harbor their own DNA (mtDNA) in multiple copies per cell and represent another target of Dox known to intercalate into DNA and to inhibit various topoisomerases. In mammalian cells, mitochondria constantly undergo fusion and fission events necessary for the sensitive regulation of cell homeostasis and rapid response upon cellular injuries (Kondadi & Reichert, 2024). Thus, morphological changes towards fission into single mitochondria has been associated with an increasing susceptibility of cells for apoptosis (Karbowski & Youle, 2003; Landes et al., 2010). Along with morphological changes, previous studies have shown dysfunction of mitochondria, such as dissipation of mitochondrial membrane potential and reduction in oxygen consumption after treatment of iPSC-derived cardiomyocytes (iCMs) with Dox (Osataphan et al., 2020). It has to be considered that the observed mitochondrial dysfunction was assessed using comparatively high doses of Dox with chronic treatment schemes. Therefore, it remains unclear whether the mitochondrial damage was caused by a direct effect of Dox or by the accumulation of multiple damages derived from nuclear DNA alterations. Here, we show that even low-dose pulse-treatment of iPSCs with Dox was sufficient to induce a shift of mitochondrial morphology towards fragmentation. It is worth to note, that mitochondrial morphology differs between iPSC cell lines depending on the genetic background of somatic origin (Harvey et al., 2016). According to previous studies, iPSC line iPS (IMR90)-4 showed mostly tubular mitochondrial phenotype with global distribution of mitochondria (Varum et al., 2011). Besides morphology, mitochondrial dynamics and mitochondrial membrane potential are strongly linked to the energetic state of the cell (Bernard et al., 2007; Sauvanet et al., 2010). Although iPSCs mainly depend on glycolysis as major pathway of energy conversion (Folmes et al., 2011; Prigione et al., 2011), mitochondrial respiration can still be evaluated. Using live-cell respirometry, we clearly demonstrated that low Dox concentrations led to a decrease in mitochondrial oxygen consumption rates and in parallel caused a loss of mitochondrial membrane potential in iPSCs. The formation of crista via invagination of the inner mitochondrial membrane into the matrix space is essential for providing an assembly platform for OXPHOS complexes (Kondadi et al., 2020; Zick et al., 2009). Since chronic exposure of cells with Dox is reported to be accompanied with disruption of mitochondrial ultrastructure (Yin et al., 2018), we further elucidated the influence of acute Dox treatment on the ultrastructure of mitochondria in iPSCs. Contrary to our expectations, pulse-treatment had no observable effect on mitochondrial ultrastructure as well as OXPHOS complex assembly and in-gel activity. Nevertheless, it is worth to mention that we showed for the first time the presence of OXPHOS supercomplexes in iPSCs. Dox is able to cross both mitochondrial membranes and to intercalate into the 16.5 kb-sized mitochondrial DNA (mtDNA) encoding 13 essential OXPHOS complex subunits (N. Ashley & Poulton, 2009; Taanman, 1999). Compared to nuclear DNA, mtDNA is not surrounded by histones protecting DNA against possible damage. Interestingly, it has been shown that upon chronic exposure to Dox, nucleoids start to condensate (Neil Ashley & Poulton, 2009). This was suggested to rather serve as a protective way to reduce intercalation of Dox into mtDNA under chronic exposure condition than to be a consequence of Dox mtDNA intercalation. Our study shows that even a mild and transient treatment with Dox is accompanied with a remodeling of mtDNA nucleoids, which apparently occurred and persisted for 48 hours after treatment. This imminent change in nucleoid size and number rather argue for a direct effect of Dox on mtDNA organization. This short time presumably also explains why we did not see any reduction of mtDNA. At this point we still cannot say whether mtDNA replication or the turnover rate of mtDNA was affected in any way upon Dox treatment. Further studies are needed to investigate the role of mtDNA turnover and possible effects on mtDNA replication in regulating cell fate in iPSCs. Besides adverse effects of Dox on stem cells in the organism, cardiomyocytes have been reported to be a strongly affected cell type upon Dox treatment. With the possibility to generate iPSC-derived cardiomyocytes (iCMs), the investigation of Dox-induced cardiotoxicity in different differentiation stages was facilitated. Our study reveals that iCMs are less susceptible towards pulse-treatment of Dox as compared to their undifferentiated counterparts. To further elucidate the influence of Dox treatment on iCMs, we performed transcriptomic profiling. We found that Dox treatment at an iPSC stage caused upregulation in genes associated with structural changes of the extracellular matrix often linked to increased fibrosis in iCMs. In particular, the upregulation of collagen and other structure-determining genes in iCMs were shown to be associated with several cardio dysfunction (Belger et al., 2024). Additionally, transcriptomic analysis revealed a consistent adverse effect of early pulse-treatment with Dox on genes essential for cardiac function such as contractility and calcium homeostasis. This is in accordance to an earlier multi-omics study investigating the effect of chronic Dox exposure to iCMs (Holmgren et al., 2018). Overall, preconditioning at the iPSC stage was associated with reduced expression levels of genes involved in glycolytic processes indicating possible malfunction of energy conversion in iCMs. Indeed, iCMs that have not undergone maturation still inherit characteristics of fetal cardiomyocytes (Lopaschuk & Jaswal, 2010; Zhou et al., 2017). This suggests a high dependency of not fully matured iCMs on glycolysis to fulfill the rudimentary energy demand of fetal cardiomyocytes. In contrast, upon late treatment expression of genes involved in apoptotic processes was enhanced. This was in accordance with a previous study where Dox treatment induced the upregulation of death receptors in cardiomyocytes (Zhao & Zhang, 2017). Taken together, pairwise comparisons of different schemes of Dox treatment allowed us to determine a timepoint scheme-specific transcriptome “fingerprint” demonstrating that preconditioning of iPSCs causes a broad and persistent genome-wide reprogramming of transcription profiles. Interestingly, many of the pathways that are negatively affected are required for cardiac function and many of the pathways that are positively affected are promoting fibrosis. Moreover, we also included a double-treatment scheme to investigate possible hormesis effects. Here, we aimed to pre-condition iCMs by pre-treatment with moderate doses of Dox in iPSCs stage. It has been shown that mild ischemia pre-exposure of the heart was beneficial in overall cell survival upon a subsequent ischemic insult (Yellon & Downey, 2003). Our study did not show any beneficial effects with regard to contractility function of iCMs function, suggesting rather an adverse effect on cardiac function upon preconditioning of cells. In accordance to these findings, it was prominent that solely iCMs that received pre-conditioning showed an increased sensitivity against Dox emphasizing highlighting the involvement of Dox-induced transcriptomic reprogramming in mediating cardiac sensitivity.

## Conclusions

Overall, we demonstrate that even mild pulse treatment with Dox of iPSCs is associated with persistent mitochondrial dysfunction and transcriptional reprogramming that affect cardiomyocyte function after differentiation. Human induced-pluripotent stem cells (iPSC) were introduced as a sensitive cell model system for investigation of the underlying susceptible processes upon Dox exposure. Apart from the timepoint of treatment, pulse treatment of iCMs revealed changes in cardiac function and fibrosis as supported by our functional and transcriptome profiling data. Our findings highlight the importance of the exposure timing and suggest that the stem cell compartment is particularly vulnerable to genotoxic stress, with long-lasting consequences that manifest after cardiac differentiation. As Dox is one of the most commonly used chemotherapeutic agents across different age groups, the question arises whether these persistent changes are responsible for the frequently observed side effects. In the adult heart, although controversially discussed, a residual stem cell pool has been reported to be present (Beltrami et al., 2003; Pfister et al., 2005; Smith et al., 2007). Thereby, possible damage to cardiomyocytes as well as to the cardiac stem cell pool could contribute to the manifestation of the lethal side effects in patients. Future studies are needed to decipher how different types of stem cells are transcriptionally preconditioned *in-vivo* after an initial exposure to sublethal doses of Dox or other genotoxic substances and how this affects heart failure and heart regeneration.

## List of abbreviations

CSC: Cardiac stem cell
CM: Cardiomyocyte
DDR: DNA damage response
DSB: DNA double-strand break
Dox: Doxorubicin
Eto: Etoposide
CCCP: Carbonylcyanid-m-chlorphenylhydrazon
iCMs: iPSC-derived cardiomyocytes
iPSC: induced pluripotent stem cell
Mt: Mock-treated
mtDNA: Mitochondrial DNA
MTT: (3-(4,5-dimethylthiazol-2-yl)-2,5-diphenyltetrazolium bromide) tetrazolium
SRB: Sulforhodamine B

## Declarations

### Ethics approval and consent to participate

not applicable

### Consent for publication

not applicable

### Availability of data and materials

The raw RNASeq dataset generated in this study including sequence reads with quality scores in fastq format study is available on SRA (Sequence Read Archive) included in the BioProject ID: PRJNA1249226 (https://www.ncbi.nlm.nih.gov/bioproject/1249226). Results of statistical analysis as described in the methods part are shown in Supplementary Table 4 showing log fold-change and adjusted p-value for each pairwise comparison per data sheet. This study was deposited as a preprint on BioRxiv.

### Competing interests

The authors declare that they have no competing interests.

### Funding

This study was supported by the German Research Foundation (DFG) RTG 2578 Project ID 417677437/GRK2578 (GF, ASR, KKo), the DFG Grant KO 6519/1-1 (AKK), and by the Ministry for Culture and Science of the State of North-Rhine Westphalia, Germany, project CERST (Center for Alternatives to Animal Testing) file number 233-1.08.03.03-121972/131–1.08.03.03–121972.

## Authors’ contributions

MW: experimental planning, data acquisition, curation, analysis, validation, investigation, visualization; methodology; writing original draft; NBA: data curation, bioinformatic analysis and methodology; YS: experimental planning, data acquisition and analysis; KKö: NGS RNA sequencing, data acquisition, resources; AD: data analysis of cardiac beating; RG: data acquisition and analysis; AB: data acquisition and analysis electron microscopy; L Ruiz-Enjuanes: data acquisition and analysis; JT: methodology, resources and data analysis; KKo: scientific discussion, methodology, resources, editing; GF: conceptualization, scientific discussion, methodology, resources, funding acquisition, editing; AKK: experimental planning, supervision, scientific discussion, methodology, ASR: study initiation, conceptualization, supervision, resources, funding acquisition, project administration, writing and editing of manuscript. All authors discussed the results and implications and commented on the manuscript at different stages. All authors read and approved the final manuscript.

## Acknowledgements

We would like to thank Ann Kathrin Bergmann and the Core Facility Electron Microscopy (CFEM) at the Medical faculty, Heinrich Heine University Düsseldorf, Germany, for support on electron microscopy, and the Genomics and Transcriptomics Laboratory (GTL), Biological and Medical Research Center (BMFZ), Medical Faculty, Heinrich-Heine-University Düsseldorf, Germany for NGS sequencing support.

## Notes

https://www.ncbi.nlm.nih.gov/bioproject/1249226

### Competing Interest Statement

The authors have declared no competing interest.

