## Supplementary figures and images for "Preconditioning of human iPSCs with doxorubicin causes genome-wide transcriptional reprogramming in iPSC-derived cardiomyocytes linked to mitochondrial dysfunction and impaired cardiac regeneration"

### Supplementary Figure 1

#### Supplementary Figure 1

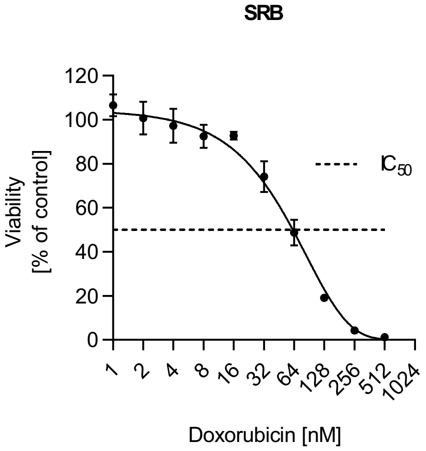

### Supplementary Figure 2

#### Supplementary Figure 2

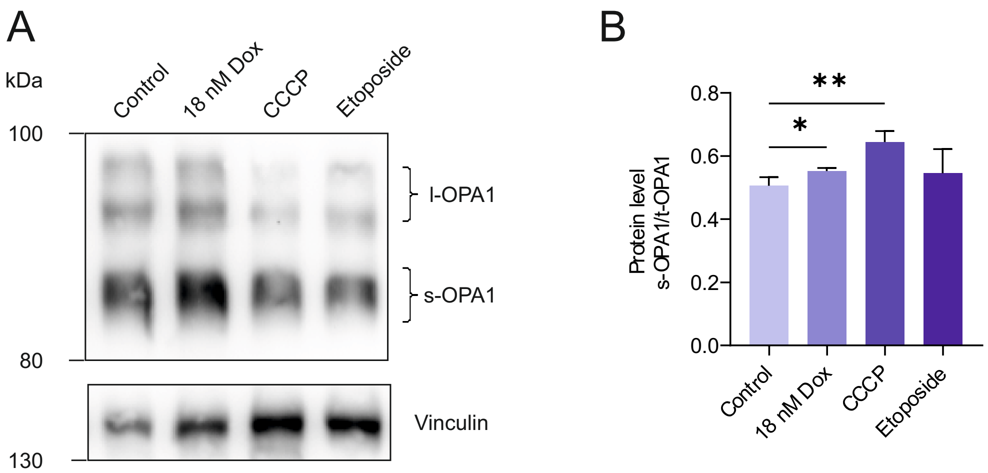

### Supplementary Figure 3

Supplementary Figure 3

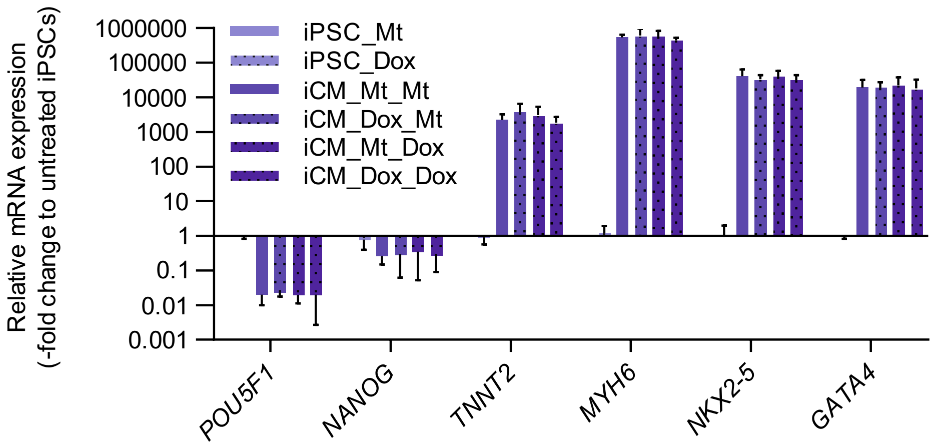

### Supplementary Figure 4

#### Supplementary Figure 4

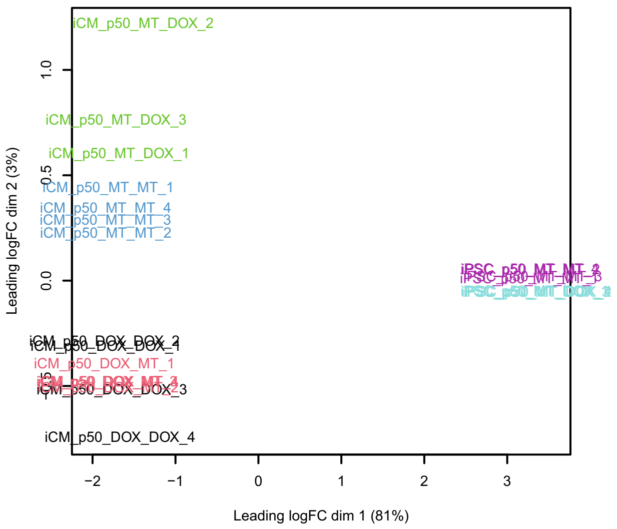

### Supplementary Figure 5

#### Supplementary Figure 5

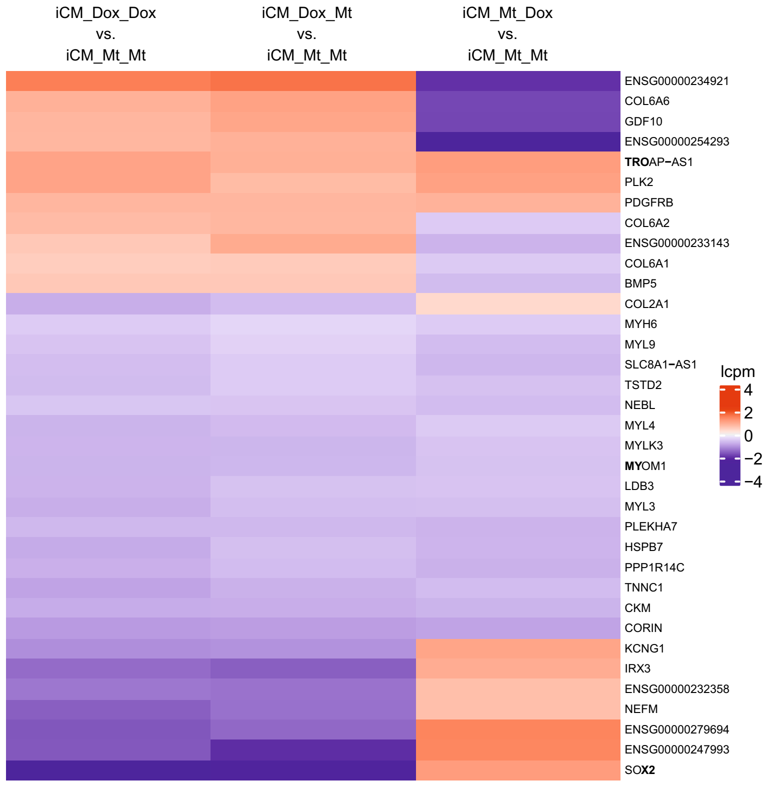

### Supplementary Figure 6

#### Supplementary Figure 6

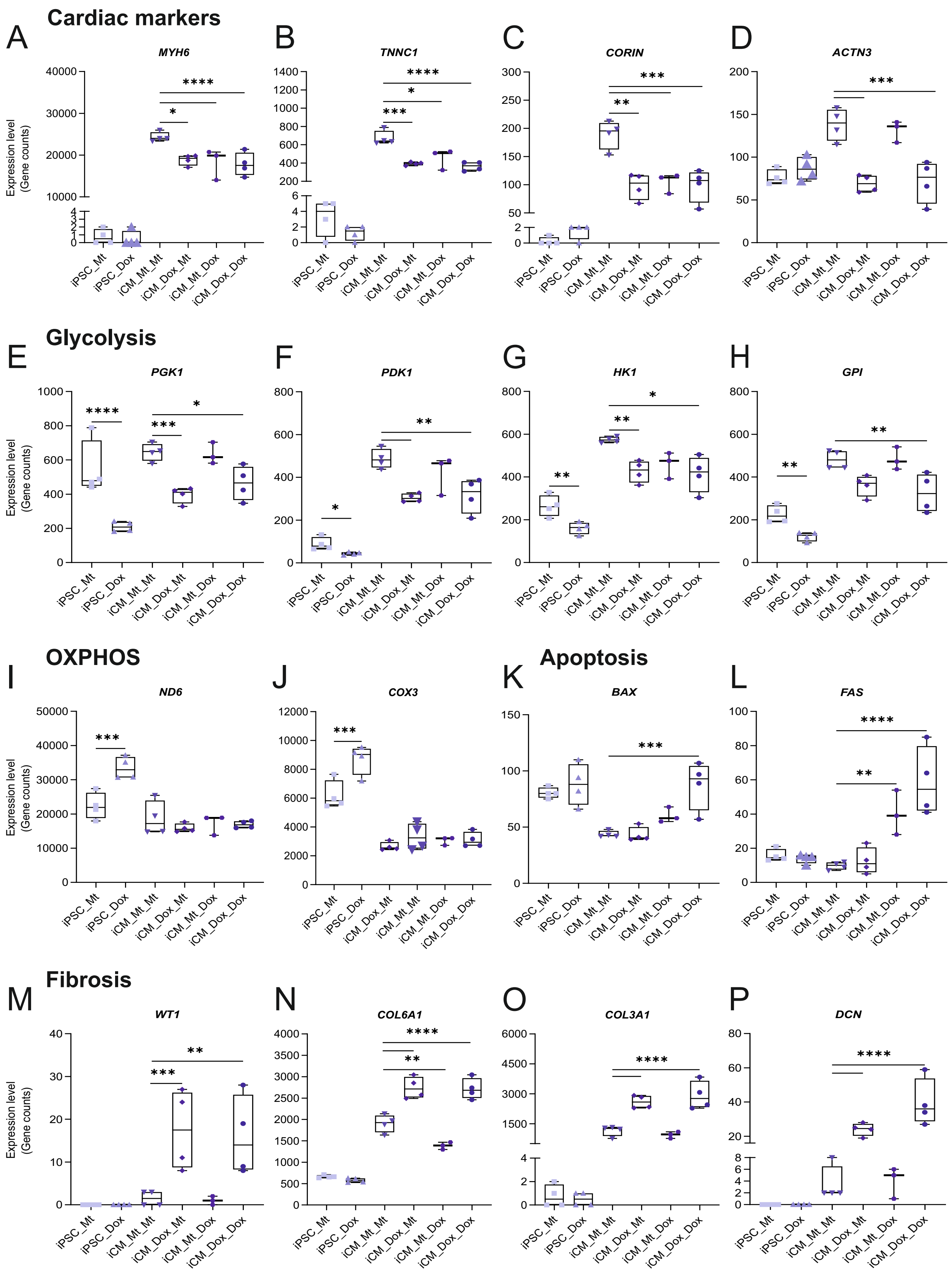

### Supplementary Figure 7

#### Supplementary Figure 7

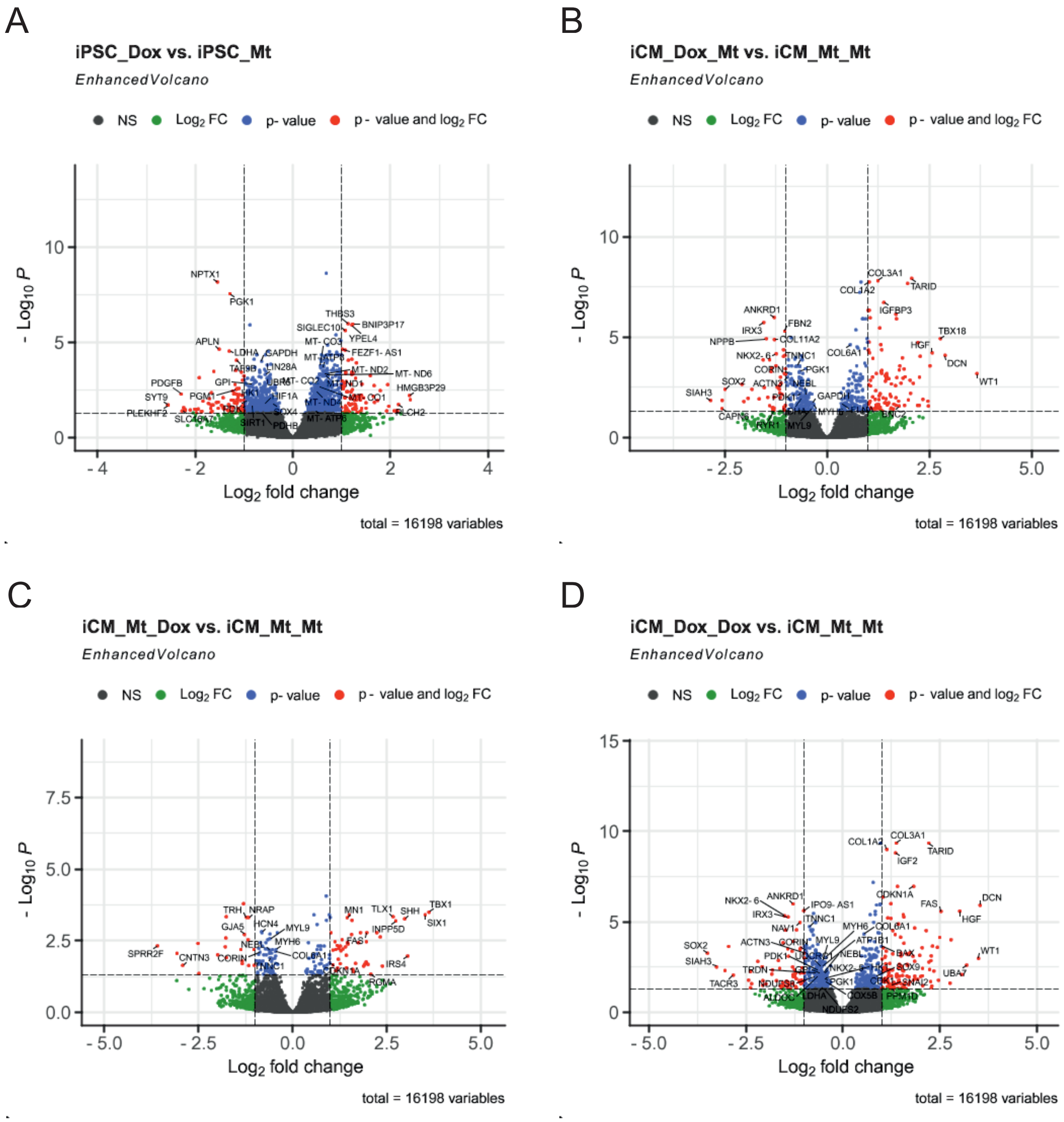

### Supplementary Figure 8

#### Supplementary Figure 8

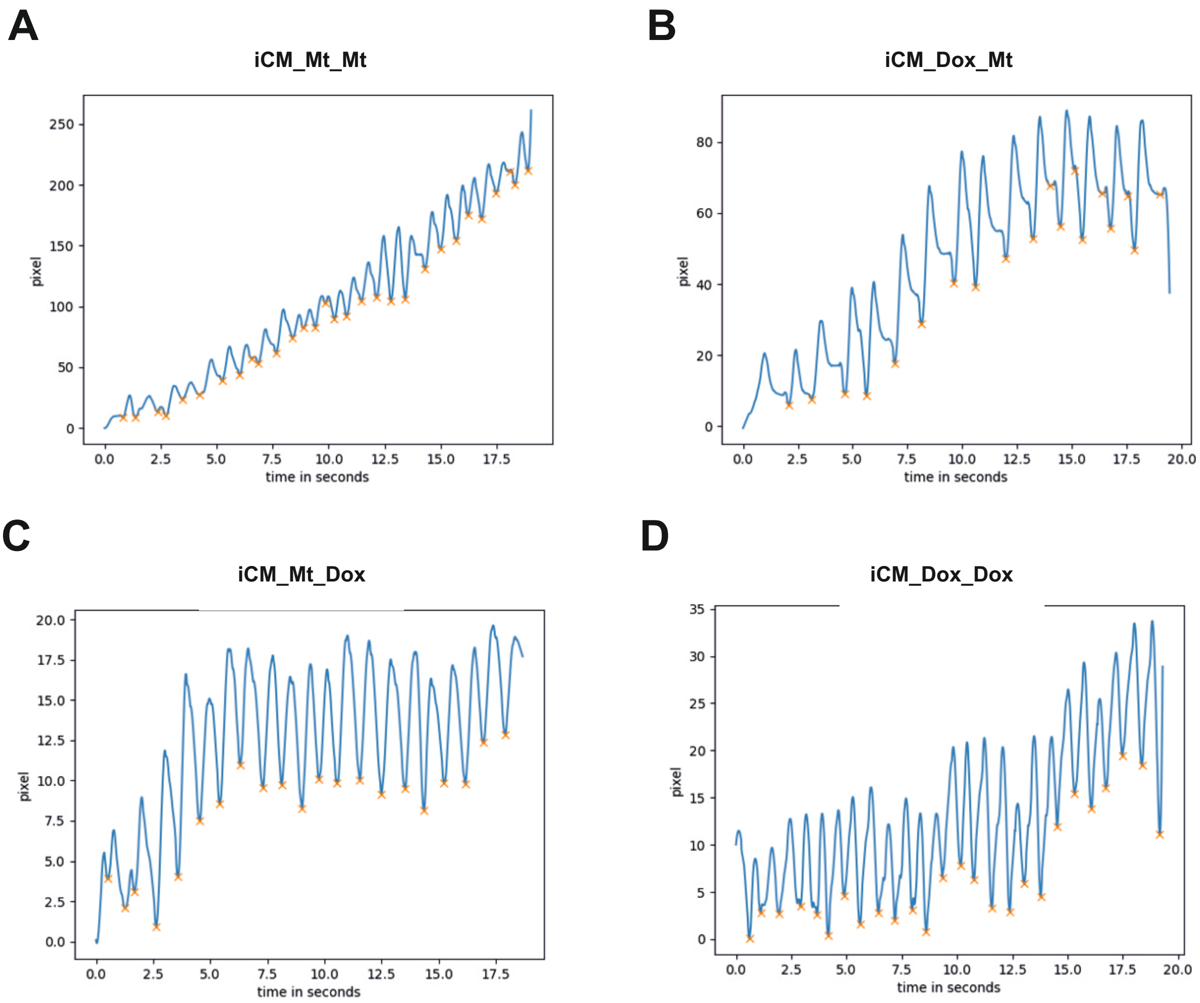
