## Supplementary materials and tables 1 to 3 and legends for "Preconditioning of human iPSCs with doxorubicin causes genome-wide transcriptional reprogramming in iPSC-derived cardiomyocytes linked to mitochondrial dysfunction and impaired cardiac regeneration"

**Supplementary Figure 1. Influence of doxorubicin pulse-treatment on viability of induced-pluripotent stem cells.** Analysis of cell viability with Sulforhodamine B assay (SRB). iPSCs were pulse-treated with diverse doses of Dox for 2 hours and cell viability was measured 48 hours after treatment with SRB assay. Mean values ± SEM of three independent biological replicates (n=3; N=8) are shown.

**Supplementary Figure 2. Influence of Dox pulse-treatment on OPA1 processing in iPSCs.** Cells were untreated (Control) or pulse-treated with 18 nM Dox for 2 hours (18 nM Dox) followed by recovery for 48 hours before total cell lysis and Western blot analysis for OPA1 isoforms was performed. As positive control for OPA1 processing, cells were treated with 10 µM Carbonylcyanid-m chlorphenylhydrazon (CCCP) for 1 hour. 10 nM etoposide for 4 hours was used as another control for DNA damage stress. The relative fraction of short OPA1 isoforms (s-OPA1) relative to all OPA1 isoforms (short and long OPA1 isoforms = total OPA1) is given. Mean values ± SD of three independent biological replicates (n=3) are shown. Student’s T-test was performed. * p-value ≤  0.05,** p-value ≤ 0.01.

**Supplementary Figure 3. Expression level of iPSC-derived cardiomyocytes treated at different timepoints of cardiac differentiation.** Relative expression levels of pluripotency markers POU Class 5 Homeobox 1 (*POU5F1*), Nanog Homeobox (*NANOG*) and cardiomyocyte marker Troponin T2 (*TNNT2*), NK2 Homeobox 5 (*NKX2-5*), Myosin Heavy Chain 6 (*MYH6*) and GATA Binding Protein 4 (*GATA4*) were measured with quantitative real-time PCR. Following samples were measured: Dox-treated iPSCs (IPSC_Dox), untreated iCMs (iCM_Mt_Mt), in iPSC stage treated iCMs (iCM_Dox_Mt), iCM treated on day7 (iCM_Mt_Dox) and iCMs treated in iPSC stage and on day 7 of cardiac differentiation (iCM_Dox_Dox). Expression levels were normalized to the untreated iPSCs control. Means ± SEM of three independent experiments (n=3; N=4) are shown.

**Supplementary Figure 4. Multidimensional scaling of all whole genome transcriptome conditions.** Results of multidimensional scaling revealing the relationship between expression profiles of the samples are shown. One biotechnical replicate was removed for the further analysis (iPSC_Mt) to avoid biased results.

**Supplementary Figure 5. Time-independent transcriptional response in iPSC-derived cardiomyocytes pulse‑treated with doxorubicin revealed partially contrary response.** Heatmap of all 35 significantly differentially regulated genes detected within the selective intersection of early treated iCMs (iCM_Dox_Mt), late treated iCMs (iCM_Mt_Dox) and double-treated iCMs (iCM_Dox_Dox)

**Supplementary Figure 6. Volcano plots of iPSCs and iCMs pulse-treated with doxorubicin compared to their untreated control.** Shown are pairwise comparisons between iPSCs and iCMs pulse-treated for 2 hours with Dox and their respective controls (iPSC_Mt and iCM_Mt_Mt, respectively). Each data point represents a gene is sorted after the mean logFC as well as the calculated p-value of four biotechnical replicate (N=3-4). In total, 16198 genes were included in this analysis. **ABCD** Volcano plot of the pairwise comparison between Dox-treated iPSCs (**A**), early treated iCMs (**B**), late treated iCMs (**C**), and double-treated iCMs (**D**) to their respective control. Top5 genes as well as genes mentioned in the result part are respectively labeled. The p-value=0.05 as well as the logFC=1 is marked with a dashed line.

**Supplementary Fig. 7. Average gene expression levels of cardiac marker genes in transcriptome conditions and their controls.** Shown are the expression levels (gene counts) of cardiac marker genes (*MYH6* (**A**), *TNNC1* (**B**), *CORIN* (**C**)*,* and *ACTN3* (**D**)), glycolytic genes (*PGK1* (**E**), *PDK1* (**F**), *HK1* (**G**), and *GPI* (**H**))**,** OXPHOS genes (*ND6* (**I**), and *COX3* (**J**)), apoptotic genes (*BAX* (**K**), and *FAS* (**L**)), and fibrosis‑related genes (*WT1* (**M**), *COL6A1* (**N**), *COL3A1* (**O**)**,** *DCN* (**P**)) resulted from RNAseq‑data of iPSCs and iCMs pulse‑treated with Dox for 2 hours with Dox and their respective controls. Each data point represents the expression level of one biotechnical replicate (N=3‑4).

**Supplementary Fig. 8.** **Visualization of peak analysis of one exemplary reference point in live‑cell videos of iCMs.** Briefly, analysis was performed with the software CardioVision. During the process, reference points were placed over the video, with a distance of 20 pixels between each point. Each reference point was analyzed and is depicted in the shown graphs. Here, peaks that were marked with an orange X and are counted as beat. **ABCD** Representative graphs of peak analysis are shown for mock-treated iCMs (iCM_Mt_Mt, **A**), iCMs already treated in iPSC stage (iCM_Dox_Mt, **B**), iCMs treated as terminal differentiated cardiomyocytes (iCM_Mt_Dox, **C**), and iCMs receiving Dox treatment in iPSC stage and iCM stage (iCM_Dox_Dox, **D**).

**Supplementary Movies 1 to 4.** Representative live‑cell videos of iPSCs under different treatment conditions. Videos were acquired at Leica DM IL LED Fluo Cellfactory with 60 frames/second. Representative videos of peak analysis are shown for mock-treated iCMs (iCM_Mt_Mt, **Movie 1**), iCMs already treated in iPSC stage (iCM_Dox_Mt, **Movie 2**), iCMs treated as terminal differentiated cardiomyocytes (iCM_Mt_Dox, **Movie 3**), and iCMs receiving Dox treatment in iPSC stage and iCM stage (iCM_Dox_Dox, **Movie 4**).

**Supplementary Table 1.** Primer sequences used for qualitative real-time PCR approaches.

| **Primer** | **Forward primer** | **Reverse primer** |
| --- | --- | --- |
| *GAPDH* | CGTAGCTCAGGCCTCAAGAC | GCTGCGGGCTCAATTTATAG |
| *HPRT1* | CCTGGCGTCGTGATTAGTG | TGAGGAATAAACACCCTTTCCA |
| *POU5F1* | CAGTGCCCGAAACCCACAC | GGAGACCCAGCAGCCTCAAA |
| *NANOG* | CAGAAGGCCTCAGCACCTAC | ATTGTTCCAGGTCTGGTTGC |
| *SOX2* | GCCGAGTGGAAACTTTTGTCG | GCAGCGTGTACTTATCCTTCTT |
| *TNNT2* | TTCACCAAAGATCTGCTCCTCGCT | AACATAAATACGGGTGGGTGCGTG |
| *NKX2-5* | AAGTGTGCGTCTGCCTTTCCCG | TTGTCCGCCTCTGTCTTCTCCA |
| *GATA4* | GCGGTGCTTCCAGCAACTCCA | GACATCGCACTGACTGAGAACG |
| *MYH6* | GGAAGACAAGGTCAACAGCCTG | TCCAGTTTCCGCTTTGCTCGCT |
| *RNR2* | AACGATTAAAGTCCTACGTGATC | TCCTTTCGTACAGGGAGGAAT |
| *MtND1* | CCACCCTTATCACAACACAAGA | GGTTCGGTTGGTCTCTGCTA |

**Supplementary Table 2. Number of significantly differentially regulated genes in the indicated pairwise comparisons.**

Number of significantly differentially expressed genes (DEGs) in Dox-treated iPSCs and iCMs versus untreated controls are shown. DEGs are categorized as upregulated, downregulated, or total, based on pairwise comparisons between treated samples and their respective controls. A total of 16,198 genes were analyzed using the voom-limma method, applying the Benjamini-Hochberg correction for multiple testing (adjusted p-value < 0.05). No log fold-change (logFC) threshold was applied. Furthermore, a downstream analysis comparing the significant DEGs resulting from iPSC_Dox vs. iPSC‑Mt comparison and the significant DEGs resulting from iCM_Dox_Mt vs. iCM_Mt_Mt resulted in the total sum 1448 DEGs influenced upon Dox treatment, among them 842 DEGs were upregulated and 606 DEGs were downregulated.

| Total number of tested genes (16198) | **iPSC_Dox vs. iPSC_Mt** | **iCM_Dox_Mt**  **vs. iCM_Mt_Mt** | **iCM_Mt_Dox**  **vs. iCM_Mt_Mt** | **iCM_Dox_Dox**  **vs. iCM_Mt_Mt** |
| --- | --- | --- | --- | --- |
| **Upregulated** | **753** | **233** | **97** | **326** |
| **Downregulated** | **519** | **193** | **89** | **306** |
| **Sum** | **1272** | **426** | **186** | **632** |

**Supplementary Table 3. Number of shared significantly differentially regulated genes in the indicated multi-comparisons.**

Number of shared significantly differentially expressed genes (DEGs) in iCMs each compared to the untreated control are shown. DEGs are categorized as upregulated, downregulated, or total, based on pairwise comparisons between different treatment conditions. Only significantly DEGs identified in a prior comparison between each treatment regimen and its respective control were included. A total of 16,198 genes were analyzed using the voom-limma method, applying the Benjamini-Hochberg correction for multiple testing (adjusted p-value < 0.05). No log fold-change (logFC) threshold was applied.

| Total number of tested genes (16198) | **DEGs specifically**  **shared by**  **iCM_Dox_Dox vs.**  **iCM Dox_Mt** | **DEGs specifically**  **shared by**  **iCM_Dox_Dox vs.**  **iCM_Mt_Dox** | **Shared by all treatment conditions** | **DEGs specific for**  **iCM_Dox_Dox** | |
| --- | --- | --- | --- | --- | --- |
| **Upregulated** | 139 | 12 | 3 | | 164 |
| **Downregulated** | 133 | 12 | 16 | | 136 |
| **Contrary regulated** | - | 1 | 16 | | - |
| **Sum** | 272 | 25 | 35 | | 300 |

**Supplementary Table 4. Excel table including information of log fold-change and adjusted p-value for each gene transcript ID, short description and gene symbol.** Each data sheet contains all 16198 genes and the respective statistics for its corresponding pairwise comparison. Significant, up- and downregulated genes can be filtered by colors of the columns (pre-filter: significant (adjusted p-value < 0.05, green), sorted descending by log fold-change (upregulation logFC > 0, blue, downregulation logFC < 0, orange).
